# Transcriptionally Active HIV Reservoirs Enriched for Interferon-Inducible APOBEC3-Related Mutational Signatures Associate with Reduced Neuroaxonal Integrity

**DOI:** 10.64898/2026.06.02.729658

**Authors:** Kazuo Suzuki, Angelique Levert, Emma Yoo, Masakazu Matsuda, Takaomi Ishida, Lucette A Cysique, John Zaunders, Ira W Deveson, Kirston Barton, Hirotaka Ode, Yasumasa Iwatani, Bruce J Brew

**Affiliations:** St Vincent’s Centre for Applied Medical Research, NSW State Reference laboratory for HIV, Sydney; St Vincent’s Clinical School, Faculty of Medicine, Sydney, New South Wales; Clinical Research Center, National Hospital Organization Nagoya Medical Center, Nagoya, Aichi, Japan; Denka Co. Ltd, Tokyo, Japan; UNSW Psychology, Sydney, NSW, Australia; Departments of Neurology and Immunology and Peter Duncan Neurosciences Unit, St Vincent’s Hospital, University of New South Wales and University of Notre Dame Sydney; Genomics and Inherited Disease Program, Garvan Institute of Medical Research, Sydney, NSW, Australia; Faculty of Medicine and Health, NHMRC Clinical Trials Centre, The University of Sydney, NSW, Australia; Department of Microbiology and Immunology, Hamamatsu University School of Medicine, University Graduate School of Medicine, Hamamatsu, Shizuoka, Japan

**Keywords:** Cell-associated HIV transcripts, Viral reservoir, APOBEC3, neuroaxonal integrity

## Abstract

Despite sustained viral suppression with effective antiretroviral therapy (ART), impaired neuroaxonal integrity persist in a subset of people with HIV-1 (PWH). We investigated whether interferon (IFN)-inducible APOBEC3-associated mutational signatures, reflected by premature stop codons (PSCs), and reservoir-associated drug resistance mutations (RS-DRMs) within transcriptionally active HIV-1 reservoirs influence neuroaxonal integrity in fully virally suppressed individuals. Peripheral CD4⁺ T cells from individuals with undetectable plasma HIV-1 RNA (n = 27) were analyzed for long HIV-1 gag/pol transcripts (>4.2 kb), and frontal white matter (FWM) neuroaxonal integrity was assessed using proton magnetic resonance spectroscopy (¹H-MRS) measurement of N-acetylaspartate (NAA).

Short HIV-1 RNA transcripts, reflecting promoter-associated transcriptional activity, were detected in all participants and were associated with reduced NAA levels in FWM, consistent with a relationship between ongoing reservoir activity and impaired neuroaxonal integrity despite virological suppression.

Long *gag/pol* transcripts (>4.2 kb) were detected in 78% of participants and exhibited marked heterogeneity in APOBEC3-associated PSC burden. PSC-containing long transcripts were associated with higher NAA, whereas long transcripts lacking such inactivating mutations were associated with reduced NAA, suggesting that APOBEC3-associated restriction of viral translational competence may contribute to differences in the downstream neurobiological effects of transcriptionally active reservoir states.

RA-DRMs were detected in 43% of participants, including triple-class resistance consistent with archived treatment exposure over more than a decade. Matching *gag/pol* sequences from reactivated virions confirmed latent reservoir origin; however, RS-DRMs showed no independent association with neuroaxonal outcomes.

Across systemic inflammatory markers, vascular risk factors, cerebrospinal fluid immune activation measures, and clinical neurocognitive outcomes (including HAND status), no consistent associations with NAA were observed.

Overall, reduced neuroaxonal integrity in the FWM was associated with the interplay between short HIV-1 RNA transcriptional activity and APOBEC3-edited translational states of long HIV-1 RNA transcripts within peripheral reservoirs, suggesting that interferon-inducible APOBEC3-associated mutational signatures may reflect biologically distinct reservoir states that correspond to neurobiological effects during suppressive antiretroviral therapy.

**Graphical Abstract:** **Interferon-Inducible APOBEC3-Associated Premature Stop Codon Signatures in HIV-1 Reservoirs and Preservation of Neuroaxonal Integrity**

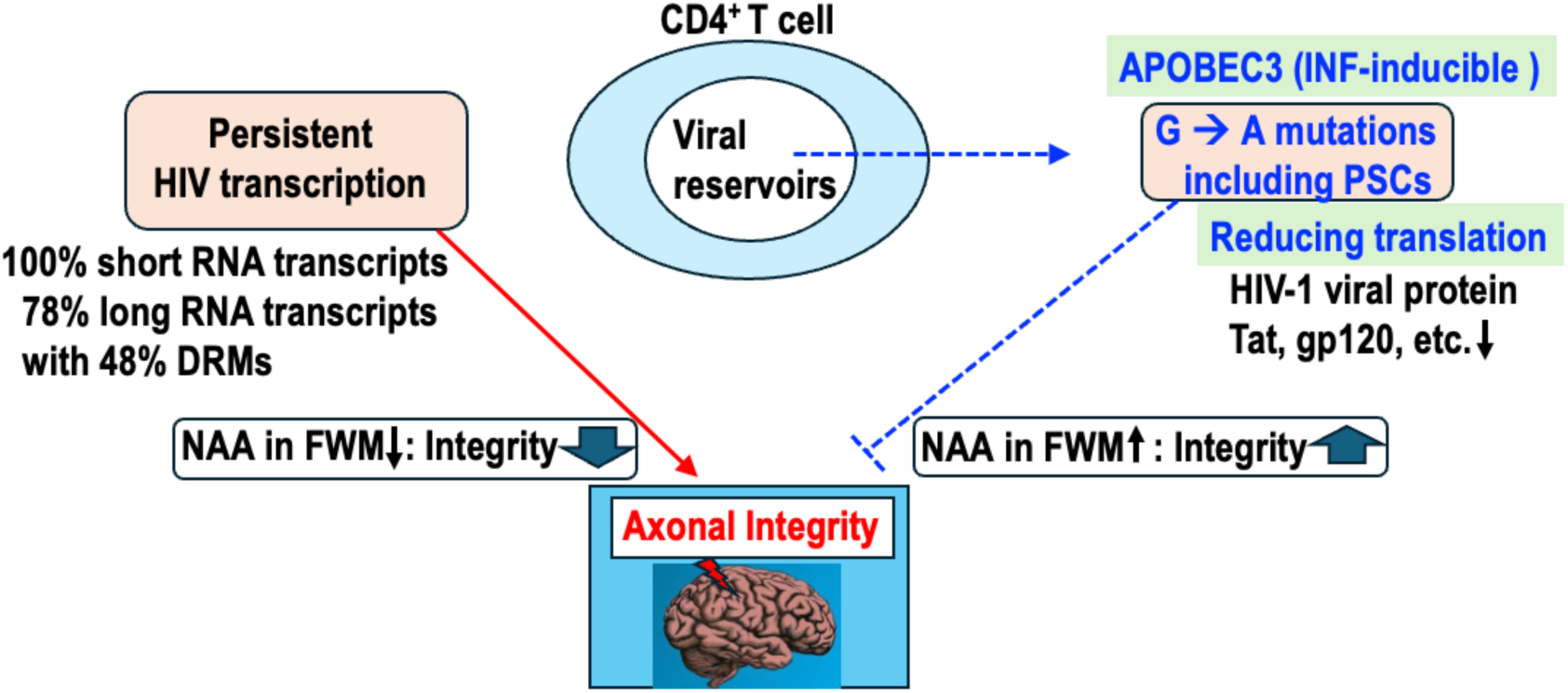

**Highlights:**

- Short HIV-1 RNA transcription within peripheral CD4⁺ T-cell reservoirs was associated with reduced frontal white matter N-acetylaspartate during suppressive ART.
- Long HIV-1 *gag/pol* RNA transcripts exhibited marked heterogeneity in interferon-inducible APOBEC3-associated G-to-A hypermutation, including premature stop codon signatures.
- Archived reservoir-associated drug resistance mutations were identified within transcriptionally active long HIV-1 reservoir-derived RNA during suppressive ART.
- Reduced neuroaxonal integrity was associated with the interplay between short HIV-1 transcriptional activity and interferon-inducible APOBEC3-associated editing of long HIV-1 transcripts with reduced predicted translational competence.

## Introduction

Despite effective antiretroviral therapy (ART) suppressing plasma HIV-1 RNA to undetectable levels, reduced neuroaxonal integrity persists in a subset of people with HIV-1 (PWH). This neuroaxonal integrity can be assessed using proton magnetic resonance spectroscopy (¹H-MRS), which reveals reduced N-acetylaspartate (NAA) in frontal white matter (FWM) ^1^. The risk of persistent neuroaxonal impairment increases with aging and comorbidities^2,3^.

The mechanisms underlying reduced neuroaxonal integrity in virally suppressed PWH remain poorly understood. Proposed contributors include neurological and psychiatric comorbidities, aging-related pathology, legacy effects of suboptimal pre-ART treatment, limited central nervous system (CNS) penetration of ART, low-level viral replication below current detection limits, and potential ART-related neurotoxicity ^4–7^. Limited ART penetration in the brain is particularly implicated, as studies reveal heterogeneity and compartmentalization of HIV-1 reservoirs across the CNS, with frontal white matter (FWM) emerging as a potential sanctuary site ^3,8,9^.

Accumulating evidence indicates that HIV-1 transcription persists despite durable plasma viral suppression. Next-generation sequencing (NGS) studies show that most HIV-1 DNA in reservoirs is replication- incompetent ^10,11^. Approximately 10–40% of proviral sequences in HIV-1 reservoir studies harbor APOBEC3-associated G-to-A hypermutational signatures, frequently generating premature stop codons (PSCs) that disrupt viral coding capacity ^12–14^. Indeed, full-length proviral sequencing studies have shown that >95% of integrated HIV-1 proviruses in ART-treated individuals are genetically defective, with only 2–5% remaining intact ^11,15^. Nevertheless, viral rebound typically occurs within weeks of ART interruption, indicating that a small subset of reservoir cells harbors replication-competent virus ^16^.

Importantly, replication incompetence does not necessarily imply biological inactivity. Defective proviruses can continue to produce viral RNA transcripts and truncated or partially translated proteins during suppressive ART, and these viral products have been associated with chronic immune activation and impaired neuroaxonal integrity ^3,17–19^. In addition, cell-associated HIV-1 RNA and DNA may engage intracellular pattern-recognition pathways, including RIG-I (retinoic acid-inducible gene I)–like signaling, which drives type I interferon (IFN) responses and interferon-stimulated gene expression in experimental models of HIV-1 persistence, even in the absence of productive viral replication ^20–23^, These observations raise the possibility that persistent reservoir transcription may contribute to end-organ injury through both innate immune signaling and translation-competent viral protein expression.

Previous studies, including ours, have demonstrated an association between HIV-1 persistence and reduced neuroaxonal integrity, based on analyses of proviral DNA and cell-associated (CA) HIV-1 RNA in cerebrospinal fluid (CSF)-derived CD4⁺ T cells ^4,24–26^. We recently reported that elevated CA short HIV-1 RNA transcripts in CSF CD4⁺ T cells were significantly associated with reduced NAA levels in FWM, suggesting that promoter-associated HIV-1 transcription may contribute to ongoing neuroaxonal impairment despite plasma viral suppression ^27^. In parallel, digital PCR (dPCR) analysis of brain tissue from virally suppressed individuals detected *gag/pol* transcripts in 42% (5 of 12) of samples, based on amplification across multiple regions of short transcripts including TAR, LTR, pol, polyA and Tat/Rev regions ^3^. These findings suggest that both short and long HIV-1 transcripts persist in the CNS despite ART. Whether long HIV-1 transcripts differ from short transcripts in neurobiological impact—through innate immune sensing, translation-competent viral protein expression (e.g., Tat), or residual replication potential—remains unknown.

A key unresolved question is whether IFN-inducible APOBEC3-associated G-to-A mutational signatures within long HIV-1 transcripts, including PSCs, functionally restrict viral translational competence and thereby influence downstream neurobiological outcomes. Although APOBEC3-associated mutational signatures are widely recognized as markers of host antiviral restriction and, in other biological contexts, have been associated with genomic instability and therapeutic resistance ^28–30^, and reservoir-associated drug resistance mutations (RA-DRMs) have been identified in persistent proviral DNA during suppressive ART ^18,31–33^, their detection within transcriptionally active HIV-1 reservoir-derived RNA transcripts has not been systematically investigated.

In this study, we investigated circulating peripheral CD4⁺ T cells as a clinically accessible HIV-1 reservoir and applied a novel assay to detect long HIV-1 *gag/pol*-frame RNA transcripts. We examined the associations between APOBEC3-related PSC signatures, RA-DRMs, and neuroaxonal integrity measured by ¹H-MRS. Our findings support a conceptual model in which reduced neuroaxonal integrity is associated with the interplay between short HIV-1 RNA transcriptional activity and APOBEC3-edited long HIV-1 RNA transcripts predicted to have reduced translational capacity, providing insight into how host antiviral restriction mechanisms may influence persistent neurobiological outcomes during suppressive ART.

## Results

### Detection of Cell-Associated HIV-1 Short RNA Transcripts in Peripheral CD4⁺ T Cells

The demographics of the 27 individuals included in this study are presented in Table S1. Participants were, on average, 56.0 years old and predominantly white Australian men with chronic, treated HIV. The dataset of more than 50 biomarkers spanning systemic inflammation, vascular risk, neuroaxonal integrity, CSF immune activation, and clinical neurocognitive outcomes (including HAND status) is available via the link provided below (see method). The median duration of HIV infection was 23 years without major neurological or systemic comorbidities. All were on suppressive ART. The median baseline CD4^+^ T-cell count was 669 cells/µL. Ten participants (37%) had an historical diagnosis of AIDS, with a median nadir CD4^+^ T cell count of 225 cells/µL.

We isolated CD4⁺ T cells from stored PBMCs by depleting CD14^+^ monocytes, followed by negative selection of CD4⁺ T cells, achieving an average purity >95% (see Methods for details). We then quantified CA HIV-1 short RNA transcripts in these CD4⁺ T cells. Short HIV-1 RNA transcripts, reflecting HIV-1 promoter (LTR)-associated transcriptional activity, were detected in all 27 individuals, with a median of 4,405 copies per 10⁶ CD4⁺ T cells (Figure 1A, C). In contrast, plasma HIV-1 RNA was detectable in only 14.8% of individuals, with a median below detection limit (Figure 1A, B). These findings are consistent with our previous reports ^27,34–37^, which demonstrated high levels of short RNA transcripts within viral reservoir cells— indicating persistent reservoir-associated transcriptional activity despite long-term plasma viral suppression.

**Figure 1.**
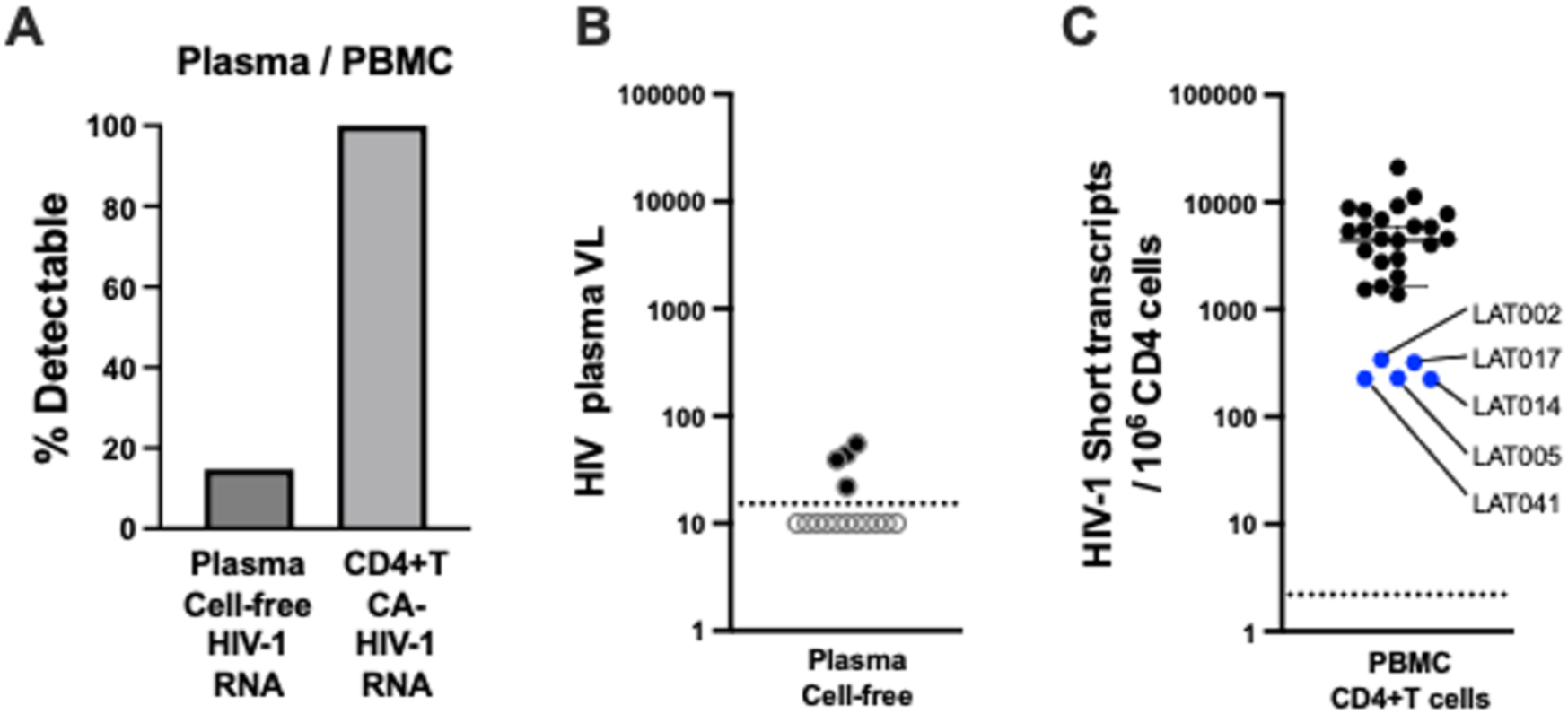
Detection of short HIV-1 RNA transcripts in CD4⁺ T cells from PWH with suppressed plasma viral load. **(A)** Comparison of detectable plasma HIV-1 RNA and short HIV-1 RNA transcript levels. Plasma viral load was undetectable in the majority of participants (14.8% had detectable levels). In contrast, short HIV-1 RNA transcripts were detected in all 27 individuals. **(B)** Plasma HIV-1 RNA levels across the cohort. The dotted line indicates the assay’s limit of detection (LOD = 20 copies/mL). Samples below the LOD were plotted at 10 copies/mL using open circles to indicate censored values. **(C)** Short HIV-1 RNA transcript levels in isolated CD4⁺ T cells, expressed as copies per 10⁶ CD4⁺ T cells (median: 4,405 copies). The dotted line represents the LOD (2 copies/10⁶ CD4⁺ T cells ^27^). Individuals highlighted in blue, with short RNA transcript levels <500 copies/10⁶ CD4⁺ T cells, failed to amplify long RNA transcripts (see Supplementary Figure 1).

### Persistent Reservoir-Associated Drug Resistance Mutations in Long HIV-1 gag/pol Transcripts

We developed a single-step RT-PCR assay targeting the HIV-1 *gag/pol* region (>4.2 kb) (see Methods) and achieved robust amplification from plasma samples (n = 85; 261 to >1,000,000 copies/mL). Long-read nanopore sequencing was performed using Oxford Nanopore Technology (ONT). Validation of this approach, demonstrating ≥95% concordance with Sanger sequencing across all DRM classes and complete agreement for major PR/RT and IN mutations, has been reported in a companion parallel study examining cardiovascular disease (CVD) associated with transcriptionally active HIV-1 reservoirs, also submitted to bioRxiv (Suzuki et al., submitted), and is consistent with prior reports of high sequencing accuracy readings with duplex sequencing analysis ^38–41^. Furthermore, we applied the HIV-1 *gag/pol* analysis to CA HIV-1 RNA derived from viral reservoirs. We identified reservoir-associated multiclass DRMs in transcripts from individuals in the CVD cohort (n = 36), all of whom had undetectable plasma HIV-1 RNA levels. Long transcripts were detected in 67% (24/36), and 50% (12/24) of these harbored RA-DRMs, indicating persistence of resistance within transcriptionally active reservoirs despite plasma suppression.

We next extended this analytic approach to the present cohort to CA HIV-1 long *gag/pol* transcripts to assess reservoir-associated drug resistance mutations (RA-DRMs). The long transcripts were successfully amplified in 21 of 27 individuals (78%) (Supplementary Figure 1). In the individuals where long transcripts were not detected, lower levels of short transcripts were observed (e.g., LAT017, LAT041, LAT014, LAT002, LAT005: blue-colored cases in Figure 1C and highlighted in light yellow in Supplementary Figure 1). This suggests a potential relationship between short and long transcript levels, suggesting a relationship between short and long transcript abundance that was further explored in subsequent ¹H-MRS analyses.

Next, we performed nanopore-based sequencing to assess the presence of RA-DRMs and APOBEC3-associated PSCs within the pol region of the 21 samples with detectable long transcripts (Supplementary Figure 1). Full-length duplex reads were analyzed using a 5% variant frequency cutoff to capture potential low-frequency DRMs in the CA viral mRNA (see Methods). PSCs were identified at variable frequencies across samples, and some samples contained *pol* sequences bearing DRMs but no PSCs. Detected DRMs within sequences lacking PSCs were categorized into three classes: protease inhibitors (PIs), non-nucleoside reverse transcriptase inhibitors (NNRTIs), and nucleoside reverse transcriptase inhibitors (NRTIs), with mutation prevalence summarized in Supplementary Figure 1 using color-coded annotations. Resistance mutations were identified in 9 of the 21 individuals: (i) Single-class resistance – NNRTI (1 case), NRTI (2 cases); (ii) Dual-class resistance – PI/NNRTI (2 cases), NNRTI/NRTI (1 case); (iii) Triple-class resistance – PI/NNRTI/NRTI (3 cases; see Supplementary Figure 1).

### RA-DRMs Reflect Both Historical and Ongoing ART Exposure

We next assessed whether RA-DRMs were associated with current and prior ART exposure. Participant LAT027, who exhibited triple-class resistance, was initially examined to investigate the relationship between reservoir-associated resistance patterns and ART history. The ART history and corresponding “Analysis point” are shown in Figure 2.

**Figure 2.**
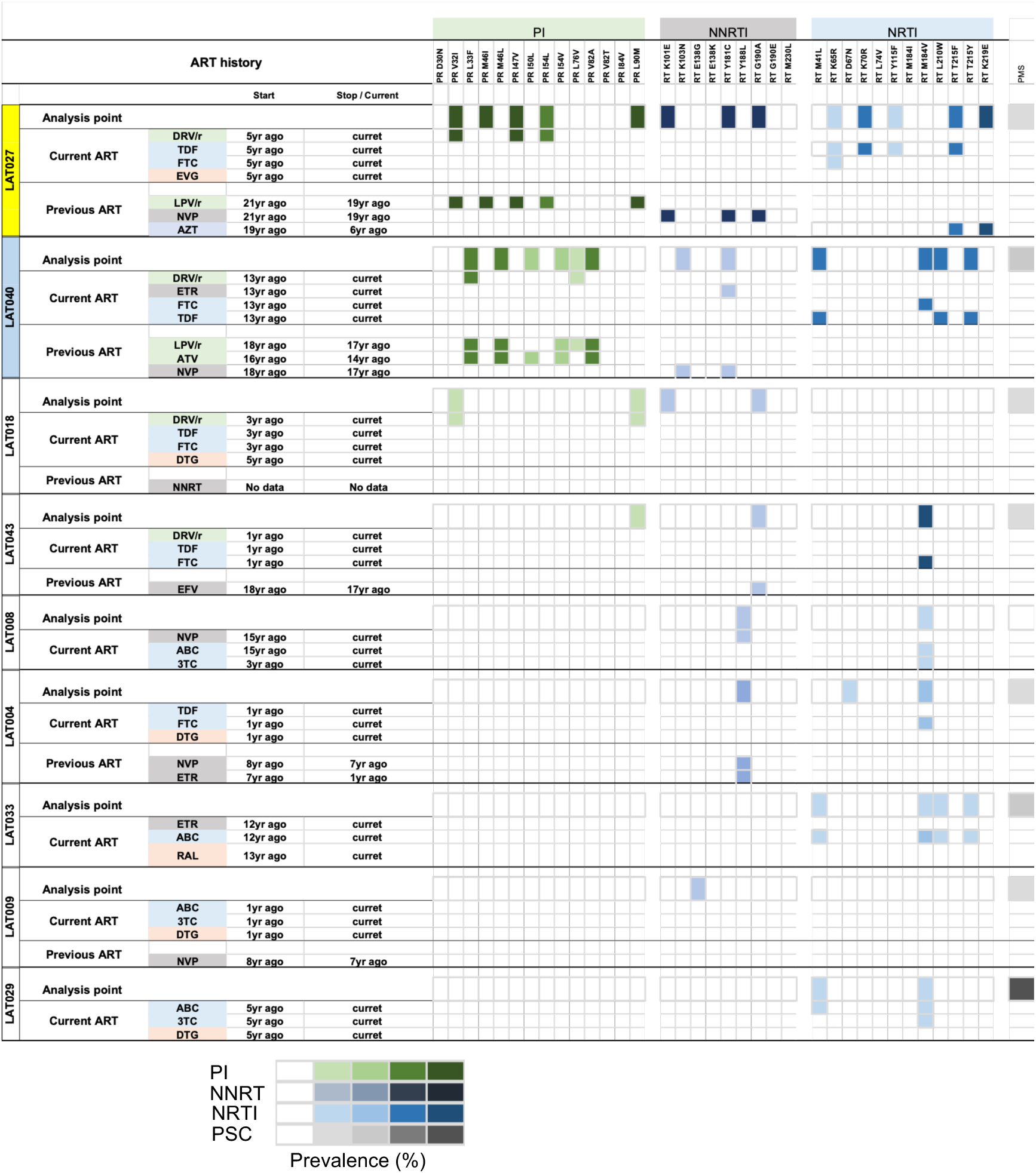
Hidden Reservoir DRMs Reflect Current and Historical ART Use. Nine individuals with the Reservoir DRMs from long RNA transcripts are shown by drug class (PI, NNRTI, NRTI) at the Analysis Point. Each DRM is aligned with current or past ART regimens, indicating start/stop dates or ongoing treatment. A prevalence bar below each point shows the proportion of DRMs potentially selected by current or prior ART. Note: Only previous ART relevant to resistance is shown.

RA-DRMs affecting PI and NRTI classes were consistent with the current ART regimen (as listed in the “Current ART” column; highlighted in yellow in Figure 2). In contrast, no integrase strand transfer inhibitor (INSTI)-associated RA-DRMs were detected despite current INSTI exposure; therefore, no INSTI-associated resistance annotations are included in Figure 2.

RA-DRMs in the PI, NNRTI, and NRTI classes were also consistent with previous treatment histories (Figure 2). Specifically, all PI- and NNRTI-associated RA-DRMs corresponded to ART regimens that had been initiated more than 21 years earlier, maintained for approximately two years, and discontinued 19 years before the analysis. Similarly, a subset of NRTI-associated RA-DRMs was consistent with zidovudine (AZT) exposure initiated more than 19 years earlier, maintained for approximately 13 years, and discontinued 6 years before the analysis. These observations demonstrate that reservoir-associated resistance signatures can remain detectable long after the corresponding therapies have been withdrawn, supporting the long-term archival of ART-associated selective pressures within persistent viral.

Furthermore, nanopore sequencing linkage analysis demonstrated that RA-DRMs associated with both historical and current ART exposure coexisted within individual viral genomes across two major viral populations, representing 53% and 11% of sequenced variants, respectively (highlighted in yellow; Supplementary Figure 2). These observations indicate that linked resistance signatures reflecting both historical and ongoing ART exposure can persist within reservoir-derived viral populations over extended periods.

A similar pattern was observed in participant LAT040, who also exhibited triple-class resistance, with the corresponding “Analysis point” shown in Figure 2. All NRTI-associated RA-DRMs and a subset of PI- and NNRTI-associated RA-DRMs were consistent with the current ART regimen (as listed in the “Current ART” column; highlighted in light blue in Figure 2).

In addition, all PI-associated and all NNRTI-associated RA-DRMs were also consistent with previous treatment histories (Figure 2). Specifically, one set of PI-associated RA-DRMs corresponded to a PI-containing regimen that had been initiated more than 18 years before the analysis, maintained for approximately one year, and discontinued 17 years before the analysis. A second set of PI-associated RA-DRMs corresponded to a different PI-containing regimen that had been initiated more than 16 years before the analysis, maintained for approximately two years, and discontinued 14 years before the analysis. Similarly, NNRTI-associated RA-DRMs corresponded to an NNRTI-containing regimen that had been initiated more than 18 years before the analysis, maintained for approximately one year, and discontinued 17 years before the analysis.

Nanopore sequencing linkage analysis further demonstrated that RA-DRMs associated with both historical and current ART exposure coexisted within individual viral genomes in three major viral populations, representing 28%, 10%, and 6% of sequenced variants, respectively (highlighted in light blue in Supplementary Figure 2). Together with the findings in LAT027, these observations indicate that linked resistance signatures associated with both prior and ongoing ART exposure can persist within reservoir-derived viral populations for many years after treatment modification or discontinuation.

Consistent findings were observed in the remaining participants (LAT018, LAT043, LAT008, LAT004, LAT033, and LAT029). With the exception of LAT009, a subset of RA-DRMs detected at the analysis points corresponded to components of the participants’ current ART regimens (Figure 2). Linkage analysis further confirmed that multiple RA-DRMs co-occurred within individual viral genomes in participants LAT018, LAT008, LAT004, LAT033, and LAT029 (Supplementary Figure 2), supporting the persistence of linked resistance signatures within reservoir-derived viral populations.

Collectively, these findings demonstrate that RA-DRMs can persist for more than two decades within reservoir-derived viral populations while maintaining linked resistance signatures associated with both historical and ongoing ART exposure, consistent with long-term maintenance of reservoir-derived viral populations and potential clonal expansion.

### Viral Outgrowth Assay (VOA) and Phylogenetic Analysis

To assess whether replication-competent virus containing intact *gag/pol* regions could be recovered from CD4⁺ T cell reservoirs, we performed viral outgrowth assays (VOAs) in three participants (LAT015, LAT027, and LAT040). CD4⁺ T cells were successfully expanded using optimized anti-CD3/CD28/CD2 stimulation (representative microscopy images shown in Supplementary Figure 3, day 6 culture).

In participant LAT040, long HIV-1 RNA transcripts were amplified from both (i) VOA culture supernatants and (ii) cell-associated RNA (CA-RNA) derived from expanded cultured CD4⁺ T cells. These sequences were compared with long HIV-1 RNA transcripts obtained from ex vivo PBMC-derived CD4⁺ T cells at baseline and from an additional ex vivo sample collected 24 months later (LAT040*, indicated by an asterisk). Phylogenetic analysis across four datasets—baseline ex vivo CA-RNA, month-24 ex vivo CA-RNA, CA-RNA from cultured CD4⁺ T cells, and VOA-derived virions—demonstrated high genetic concordance within participant LAT040 (blue-colored labels in Figure 3), indicating preservation of closely related viral lineages across compartments and over time.

**Figure 3.**
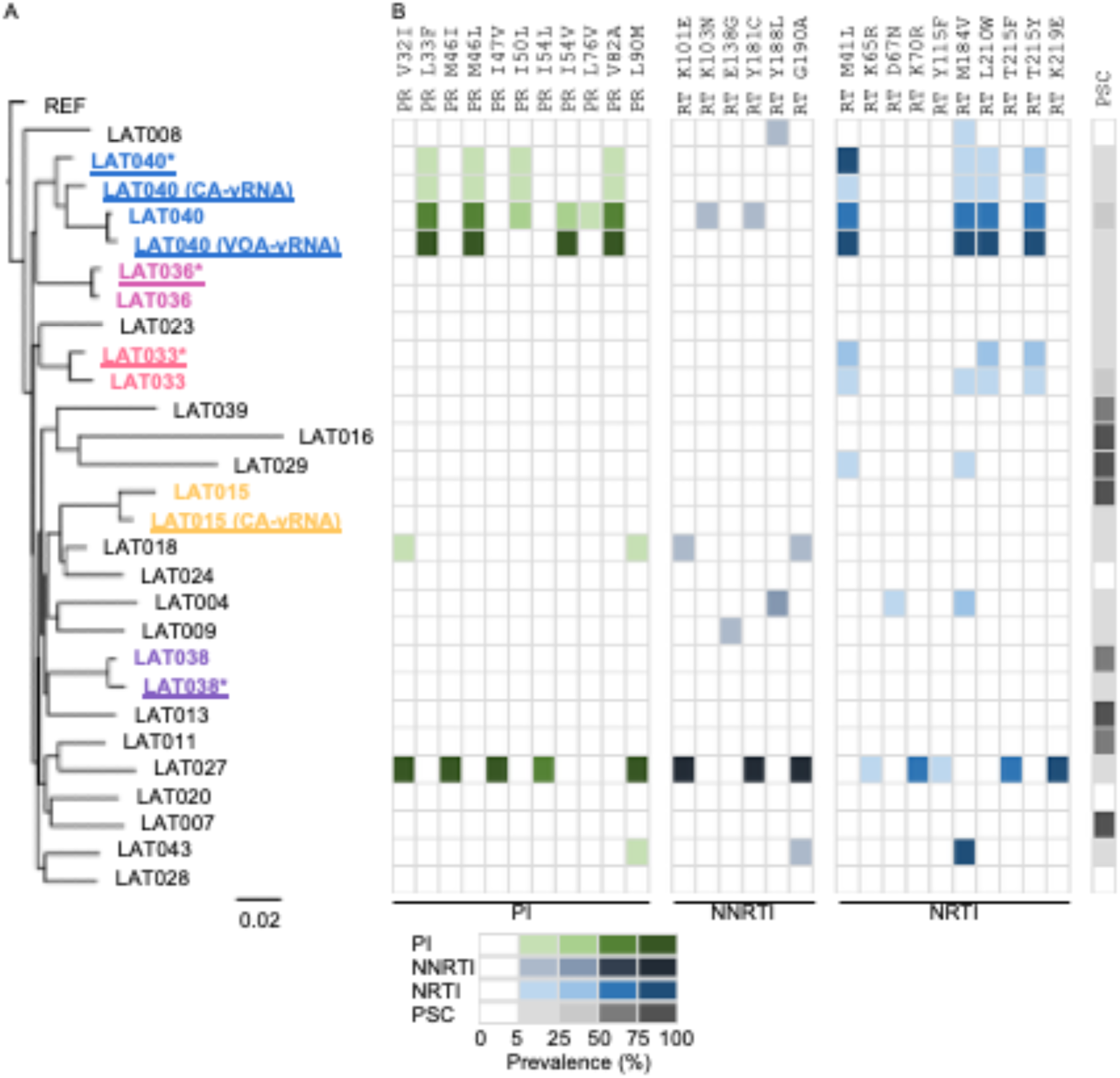
Phylogenetic relationships and major drug resistance mutations (DRMs). **(A)** Approximate maximum-likelihood tree of consensus *pol* sequences from each sample, constructed using FastTree with HXB2 (GenBank: K03455) as the reference. The sample labels are shown in the same order and manner to that in Supplementary Figure 2. Samples from the same individual—including additional CA-vRNA or VOA-vRNA sequences collected at later time points (18 or 24 months)—are color-coded and underlined; asterisks (*) denote additional samples. **(B)** Prevalence of major DRMs in *pol* sequences lacking PSCs, color-coded by sample. The proportion of sequences containing PSCs in *pol* is also shown.

VOA-derived viral RNA could not be amplified from culture supernatants in participants LAT015 and LAT027. However, CA-RNA was successfully recovered from expanded cultured CD4⁺ T cells in LAT015 (LAT015 CA-vRNA; Figure 3). In this participant, phylogenetic analysis demonstrated close genetic similarity between ex vivo CA-RNA and CA-RNA recovered from cultured cells, consistent with the persistence of related viral populations during in vitro expansion (blue-colored labels in Figure 3).

To further evaluate longitudinal stability of reservoir-derived viral populations, CA-RNA was analyzed at two time points in 3 additional participants (LAT033, LAT036, and LAT038). Across all participants examined longitudinally (including LAT040), reservoir-derived viral sequences remained highly conserved over time, with no evidence of major phylogenetic divergence. Additional longitudinal samples are identified by color-coded, underlined labels marked with asterisks in Figure 3.

Collectively, these findings demonstrate substantial genetic stability of reservoir-derived HIV-1 populations across longitudinal sampling and in vitro expansion. Furthermore, the close phylogenetic relationship between ex vivo CA-RNA, cultured-cell CA-RNA, and VOA-derived viral RNA supports the persistence of transcriptionally active viral lineages within the reservoir. These observations provide a framework for investigating post-transcriptional constraints on viral protein expression, including APOBEC-associated premature stop codon formation, as examined in the following section.

### CA HIV-1 Short and Long RNA Transcripts and Their Relationship with Neuroaxonal Integrity

We examined the association between HIV transcriptional activity in CD4⁺ T cells derived from peripheral blood and neuroaxonal integrity. Building on prior findings demonstrating a significant association between short HIV-1 RNA in CSF-derived CD4⁺ T cells and NAA levels in FWM ^27^, we observed a similar significant inverse correlation between short HIV-1 RNA transcripts in PBMC-derived CD4⁺ T cells and FWM NAA levels, as measured by ¹H-MRS (Figure 4A). A weaker but still significant inverse correlation was also observed in the posterior cingulate cortex (PCC), consistent with reduced neuroaxonal integrity (Figure 4B). In contrast, no association was observed with NAA in the caudate nucleus, a region previously shown to have high HIV burden in the pre-ART era (Figure 4C).

**Figure 4.**
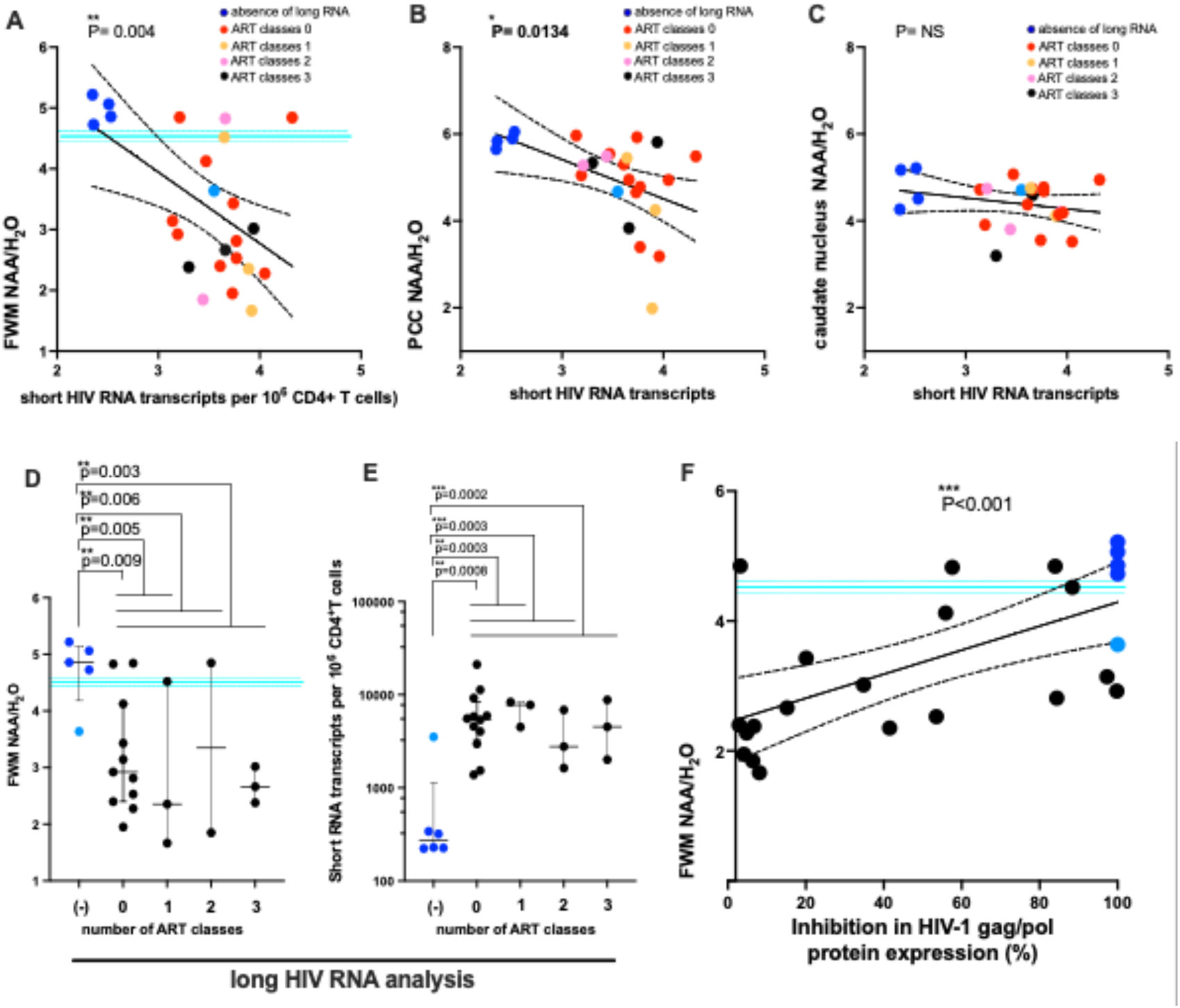
Dual impact of HIV-1 RNA transcription and translation on neuroaxonal integrity. **(A)** Inverse correlation between CA short HIV-1 RNA transcript levels and NAA levels in the FWM. **(B)** Inverse correlation between CA short HIV-1 RNA transcripts and NAA levels in PCC. **(C)** Association between CA short HIV-1 RNA transcripts and NAA levels in the caudate nucleus. **(D)** Association between FWM NAA levels and the presence or absence of DRMs in the HIV-1 reservoir. Individuals in blue correspond to those highlighted in Figure 1C. **(E)** Relationship between short HIV-1 RNA transcript levels and the presence or absence of reservoir DRMs. **(F).** Impact of APOBEC-associated PSCs, which prevent *gag/pol* translation, on neuroaxonal integrity. Six individuals failed to amplify long RNA transcripts, indicated by blue circles (4 dark blue, 1 light blue), consistent with 100% translational inhibition. *Note: NAA data for the frontal white matter (FWM), posterior cingulate cortex (PCC), and caudate nucleus were unavailable for LAT014, a dark blue–colored individual in Figure 1C who exhibited low levels of short HIV-1 RNA transcripts and for LAT021 (light blue), who lacked detectable long RNA transcripts.* *The solid blue line in panels **A, D** and **F** represents the mean normative FWM NAA level at age 56; upper and lower dotted lines correspond to ages 45 and 67, respectively ^1^.*

We next examined whether long CA-RNA transcripts were associated with neuroaxonal integrity. Individuals with detectable long transcripts had significantly lower FWM NAA levels compared to those without detectable long transcripts (Figure 4D, *p* = 0.003). In contrast, no significant associations were observed in PCC or caudate NAA (Supplementary Figures 4A, 4B).

Participants in whom long HIV-1 RNA transcripts were not detected (LAT002, LAT017, LAT005, LAT041; highlighted in blue) showed higher FWM NAA levels and lower levels of short HIV-1 RNA transcripts (Figure 1C and Figure 4E), consistent with better preservation of neuroaxonal integrity. In contrast, participants with detectable long transcripts showed reduced FWM NAA (Figure 4D). Participant LAT021, shown as a light blue point in Figure 4, exhibited detectable short HIV-1 RNA transcripts in CD4⁺ T cells (3,521 copies) and was therefore considered separately from the long-transcript-negative group. FWM NAA data were unavailable for participant LAT014. Overall, these findings suggest an association between long HIV-1 RNA transcripts and reduced neuroaxonal integrity, as well as a relationship between short transcript levels and the likelihood of detecting long transcripts in viral reservoirs (Figure 4E).

The number of ART-associated RA-DRMs within long transcripts was not associated with NAA levels, suggesting no relationship between RA-DRM burden and neuroaxonal integrity in this cohort (Figure 4D). Similarly, no association was observed between short RNA transcript levels and the number of DRMs within long transcripts (Figure 4E).

### APOBEC3-associated hypermutation and translational constraints in HIV-1 reservoir-derived RNA

Building on the presence of transcriptionally active reservoir-derived viral RNA and persistent RA-DRMs, we next investigated whether these RNA species exhibited evidence of post-transcriptional restriction affecting viral protein coding capacity.

Sequence analysis of long HIV-1 RNA transcripts revealed frequent G-to-A hypermutational patterns consistent with host APOBEC3-associated editing, resulting in the introduction of premature stop codons (PSCs) across multiple genomic regions. These APOBEC-associated mutations were observed within transcriptionally active viral RNA populations identified in CD4⁺ T cells under suppressive antiretroviral therapy.

The distribution of PSC-containing transcripts varied across participants and was enriched within subsets of reservoir-derived viral populations identified in the preceding analyses. These findings suggest that transcriptionally active viral reservoirs may include a substantial proportion of transcripts with disrupted coding potential, limiting production of full-length viral proteins.

To further explore the possible functional consequences of these mutations, we developed an exploratory model to assess the relationship between inferred *gag/pol* translational capacity and neuroaxonal integrity. In this model, reduced detection of intact *gag/pol* RNA transcripts was used as a proxy for decreased potential for full-length *gag/pol* protein expression, acknowledging that this represents an indirect estimate of inferred *gag/pol* translational competence. For modeling purposes, cases in which long gag/pol-frame (unspliced) genomic RNA was not detected were classified as having absent potential for full-length *gag/pol* translation, based on long transcript detection status and analyzed independently from PSC frequency, consistent with 100% translational inhibition within the framework of this exploratory model. Using this framework, we observed a significant association between inferred *gag/pol* translational competence in CD4⁺ T cells and neuroaxonal integrity, as measured by NAA levels in FWM as shown in Figure 4F (p < 0.001).

Notably, participants with higher levels of PSC-associated APOBEC3 mutation signatures in the *gag/pol* region exhibited higher NAA levels in FWM, whereas those with lower PSC levels—indicating a greater proportion of intact *gag/pol* transcripts—exhibited lower NAA levels (Figure 4F). In Figure 4A and 4F, the solid horizontal line indicates the mean normative FWM NAA level at age 56, with the upper and lower dotted lines representing the corresponding normative values at ages 45 and 67, respectively.

## Discussion

This study demonstrates that both short and long HIV-1 RNA transcripts are associated with reduced neuroaxonal integrity, predominantly in FWM, with no significant associations observed in deep gray matter regions. This regional specificity, also observed in prior studies of CNS reservoirs ^9^, contrasts with the classic pattern of HIV-1-associated brain injury in the pre-ART era, which predominantly affected deep gray matter, particularly the basal ganglia ^1^. Such a shift in anatomical vulnerability may reflect the changing biology of HIV-1-associated neuropathogenesis under suppressive antiretroviral therapy. Importantly, APOBEC3-associated G-to-A mutational signatures within long HIV transcripts, including PSCs, were associated with relatively preserved FWM NAA levels, suggesting that host restriction–linked mutational processes may attenuate the translational competence of transcriptionally active reservoir-derived transcripts and thereby influence downstream neuroaxonal injury.

To our knowledge, this is the first in vivo study to identify both APOBEC3-associated PSC signatures and RA-DRMs within transcriptionally active HIV-1 RNA derived from peripheral blood CD4⁺ T-cell reservoirs, and to assess these molecular signatures in relation to measures of CNS integrity. These findings provide biological evidence that long HIV-1 transcripts retaining predicted translational competence may contribute to ongoing HIV-associated neurotoxicity, whereas extensive APOBEC3-associated G-to-A hypermutation resulting in PSCs may reduce predicted translational competence and potentially attenuate downstream neurobiological effects.

Mechanistically, these findings are consistent with potential involvement of innate antiviral sensing within reservoir cells. HIV-1 RNA species have been reported to activate RNA-sensing pathways involving RIG-I, leading to type I interferon signaling, which is a known inducer of APOBEC3 expression ^20–23^. Although these pathways were not directly measured in the present study, the short HIV-1 RNA transcripts detected by the Double-R assay may represent viral RNA species capable of engaging innate immune sensing pathways. Our data are therefore consistent with a model in which elevated short HIV-1 RNA transcription increases the likelihood of generating long transcriptionally active viral RNA species while being accompanied by interferon-driven APOBEC3 activity. Under this model, long HIV-1 transcripts generated in the context of persistent short RNA transcription may become subject to APOBEC3-associated editing, resulting in transcripts enriched for premature stop codons and reduced predicted translational competence.

These observations suggest an interplay between short HIV-1 transcriptional activity and APOBEC3-edited translational states of long HIV-1 transcripts within viral reservoirs. PSC burden appears to be associated with reduced predicted translational competence. HIV-1 regulatory proteins such as Tat and Rev play essential roles in viral gene expression, with Tat required for transcriptional elongation ^42–46^ and Rev required for nuclear export of full-length and partially spliced viral RNA ^47–50^. Because the rev coding region lies immediately downstream of the sequenced pol region analyzed in this study, PSC-associated mutations within gag/pol may potentially impair expression of functionally intact Rev. Importantly, Tat has previously been implicated in HIV-associated neurotoxicity ^51–54^ while Rev is required for efficient export of singly spliced env transcripts ^55^, which encode neurotoxic viral proteins such as gp120 ^52,56,57^. Accordingly, APOBEC3-associated translational constraints may reduce efficient viral RNA export to the cytoplasm and limit production of neuroactive viral proteins, including Tat and gp120.

Furthermore, multiple PSCs within phylogenetically related transcripts support sustained persistence of reservoir-derived viral populations rather than transient transcriptional bursts, supporting a link between peripheral reservoir biology, APOBEC3-associated restriction, and CNS integrity during suppressive ART.

These findings have direct implications for HIV brain disease and cure or analytical treatment interruption studies ^32,58–64^. Current ART does not consistently reverse mild neurocognitive impairment despite plasma viral suppression. Our data suggest that ongoing HIV transcription within reservoir cells is more closely associated with neuroaxonal integrity in FWM than measures of low-level replication or ART exposure, highlighting reservoir transcription as a potential therapeutic target that may be influenced by APOBEC3 biology.

Individuals with undetectable long HIV-1 RNA transcripts and low levels of CA short HIV-1 RNA may represent potential candidates for cure-directed interventions, as they exhibited relatively preserved neuroaxonal integrity in this cohort. Accordingly, CA short and long HIV-1 RNA transcripts may serve as sensitive biomarkers of residual reservoir activity beyond standard plasma assays.

In this fully suppressed cohort, a broad panel of biomarkers spanning systemic inflammation, vascular risk, CSF immune activation, and clinical neurocognitive outcomes showed no detectable association with neuroaxonal integrity. These findings suggest that conventional systemic biomarkers may not fully reflect mechanisms underlying persistent neuroaxonal alterations in individuals with undetectable plasma HIV-1 RNA during ART. In contrast, reservoir-derived transcriptional activity, particularly short HIV-1 RNA transcripts, showed the strongest association with reduced NAA, supporting a model in which CNS integrity may be influenced by reservoir transcriptional states rather than systemic immune activation alone.

Together, these findings support a model in which neuroaxonal dysfunction during suppressive ART is associated with ongoing reservoir-derived transcription rather than systemic inflammation or classical viral replication alone. This process may be influenced by APOBEC3-associated editing, which introduces PSCs that constrain viral protein translation and may reduce neurotoxicity.

Some integrase strand transfer inhibitor (INSTI) and non-nucleoside reverse transcriptase inhibitor (NNRTI) mutations may reflect APOBEC-associated hypermutation rather than true drug-selective pressure, requiring further validation. Additionally, high PSC burden was rarely associated with substantial resistance mutation accumulation, suggesting partially distinct pathways of reservoir persistence and restriction. Longitudinal studies will clarify temporal dynamics.

More broadly, our findings support a conceptual framework in which persistent HIV reservoir transcription may contribute to chronic end-organ injury through interconnected innate immune and translational pathways during suppressive ART. In this model, ongoing short HIV-1 RNA transcription may promote interferon-associated innate immune activation, which could simultaneously induce APOBEC3-associated G-to-A hypermutation within long HIV-1 transcripts while also contributing to inflammatory processes linked to reduced neuroaxonal integrity. Although interferon signaling and APOBEC3 activity were not directly measured in this study, the observed association between APOBEC3-related PSC signatures and neuroaxonal outcomes is consistent with a shared interferon-inducible reservoir state. This framework may also help integrate our observations across both neuroaxonal and cardiovascular reservoir studies (Suzuki et al., submitted), supporting the possibility that persistent transcriptionally active HIV reservoirs contribute to chronic end-organ disease despite suppressive plasma viremia.

In summary, HIV-associated neuroaxonal alterations persist despite effective plasma viral suppression and are most closely associated with transcriptionally active reservoir-derived RNA rather than systemic immune activation alone. APOBEC3-associated editing introduces PSCs that may restrict viral protein translation, including regulatory proteins such as Tat and Rev, thereby potentially reducing neurotoxicity. Persistent reservoir-associated transcriptional activity may additionally contribute to interferon-associated innate immune and inflammatory pathways linked to chronic neuroaxonal alterations during suppressive ART. These findings identify reservoir transcriptional activity as an important contributor to CNS integrity in treated HIV infection and support targeting viral transcriptional programs alongside host restriction pathways as potential strategies for neuroprotection and cure development.

## Materials and Methods

### Participants

Twenty-seven HIV–infected male participants (Table S1) receiving fully suppressive antiretroviral therapy (ART) were enrolled in an ongoing prospective study of CNS HIV latency and NeuroHIV biomarkers (ClinicalTrials.gov Identifier: NCT02989285). Inclusion criteria required stable HIV-1 infection with sustained viral suppression. The study protocol was approved by the St. Vincent’s Hospital Human Research Ethics Committee (HREC/15/SVH/425), and all participants provided written informed consent prior to enrolment

### Isolation of CD4^+^ T cells from peripheral blood mononuclear cells (PBMC)

PBMCs were isolated from acid-citrate dextrose (ACD) anti-coagulated blood by density centrifugation using Ficoll-Paque Plus (GE Healthcare, Chicago, IL, USA). Isolated PBMCs were cryopreserved in heat-inactivated, filter-sterilized bovine serum (ThermoFisher Scientific, Waltham, MA, USA) containing 10% dimethyl sulfoxide (DMSO; Sigma-Aldrich, MO, USA) using a controlled-rate freezer (Planer, Middlesex, UK) and stored in the vapor phase of liquid nitrogen.

CD4⁺ T cells were isolated from thawed PBMCs using a previously described method^27^. Average of CD4⁺ T Cell purity was exceeded 95%, as confirmed using the same protocol.

### Short HIV RNA transcripts analysis

RNA extraction and quantification of short HIV-1 RNA transcripts were performed as previously described ^27,34^. Primers and probes targeted the highly conserved “R” region within both the 5′ and 3′ long terminal repeats (LTRs; Double-R assay), enabling detection of total spliced and unspliced transcripts by one-step reverse transcriptase PCR (RT-PCR; WO2018/045425, PCT/AU2017/050974) as a measure of HIV-1 promoter activity. Amplicons were detected using precision image analysis of πCode MicroDiscs (PlexBio) on the IntelliPlex platform, as described previously ^27,34^.

### Long HIV-1 RNA transcripts analysis

A long-range RT-PCR protocol was used for selective amplification of full-length, in-frame *gag/pol* transcripts (>4.5 kb), as outlined in the Australian provisional patent application No. 2024903454. Briefly, long RNA transcripts were amplified using a one-step RT-PCR protocol followed by nested long-range DNA PCR. This approach generated >4.5 kb amplicons of the *gag/pol* frame region for downstream sequencing.

### Nanopore Sequencing analysis

DNA library for nanopore sequencing were prepared as previously reported ^41^. Briefly, RT-PCR amplicons of *gag*/*pol* transcripts were purified using the AMPure XP beads (Beckman Coulter, Brea, CA, USA), and their concentration was measured using a Qubit 4 Fluorometer with the Qubit 1X dsDNA HS Assay Kit (Thermo Fisher Scientific).

To prepare DNA library, the amplicons were end-repaired and were subsequently ligated with native barcodes and adapters. Long Fragment Buffer was used for the final wash. The prepared DNA library was quantified using Qubit and then loaded onto an R10.4.1 flow cell (ONT). Sequencing was performed on a MinION Mk1C device using MinKNOW software (ONT), with fast base-calling enabled for a one-hour run.

### Analyses of the Drug resistance mutations and phylogenetic tree

Based on duplex reads obtained from nanopore sequencing for each sample, prevalence of drug resistance mutations (DRMs) and PSCs within pol were examined as previously described with a slight modification ^41^. In this study, the major DRMs listed in Stanford HIV database (https://hivdb.stanford.edu/) were adopted for the analyses. DRMs that could arise from APOBEC3 editing were defined as shown in the previous report ^41^. To explore the DRMs and PSCs from the reads, reads were mapped on a partial HXB2 sequence (GenBank ID: K03455, 2,253-5,096 nt) as the reference. Then, reads that fully cover sequences between the 1st codon of protease (PR) and the 264th codon of integrase (IN) (2,253-5,021 nt according to HXB2 coordinate) were selected, because all the major DRMs appear between the PR 30th codon and IN 263rd codon. Coverage for selected reads exceeded 100 per sample, consistent with prior work ^41^. Analyses focused on the region between PR codon 1 and IN codon 264. For each sample, per-site prevalence of major DRMs and PSCs was determined. Additionally, prevalence of genotypes harboring unique DRM patterns, with or without PSCs, was assessed. A 5% cutoff was applied to detect DRMs, justified as follows: (i) duplexed reads with bidirectional strand analysis; (ii) >100 reads per site, with at least 100× depth, so a 5% variant is supported by ≥5 reads; (iii) low-frequency variants (5–10%) may be clinically significant ^65^, especially in ART-experienced individuals ^66^.

Consensus nucleotide sequences were generated by selecting the dominant base at each position from the curated reads. Multiple alignments were performed using MAFFT v7.372 (https://pubmed.ncbi.nlm.nih.gov/23329690/), followed by inference of approximately-maximum-likelihood phylogenetic tree with FastTree ver. 2.1^67^. Trees were visualized with FigTree v1.4.4 (http://tree.bio.ed.ac.uk/software/figtree/).

### Proton Magnetic Resonance Spectroscopy (¹H MRS)

In this study, the term ‘neuroaxonal integrity’ refers specifically to FWM NAA measurements, whereas ‘neuronal integrity’ is used more broadly when referring to NAA measurements across multiple brain regions of PCC and caudate nucleus.

¹H MRS brain scans were analyzed as previously described ^27^. Briefly, spectra were acquired using a Philips 3T Ingenia scanner (Philips, Best, Netherlands). Cerebral metabolite concentrations were quantified using point-resolved spectroscopy (PRESS) with echo time (TE) = 40 ms and repetition time (TR) = 2000 ms. Data were collected from the FWM, posterior cingulate cortex (PCC), and caudate nucleus. Metabolites were fitted in JMRUi V4 and referenced against unsuppressed H^2^O.

As part of a comprehensive biomarker phenotyping approach, more than 50 biomarkers relevant to neuroaxonal integrity were assessed across multiple biological domains, including brain neurochemistry [NAA (primary neuroaxonal integrity marker), Cho, and mI], regional brain vulnerability [PCC, caudate, and FWM], cardiovascular and hemodynamic function [blood pressure, D:A:D cardiovascular risk scores, and Framingham risk], metabolic and cardiac injury markers, systemic inflammation [CRP, IL-6, and plasma neopterin], CSF viral protein biology, CSF immune activation, substance use disorders, cardiovascular risk factors, HIV-related clinical parameters, and hematological indices. The full dataset is available via the link provided below.

### Viral Outgrowth Assay (VOA)

VOA was performed for participants LAT015, LAT027, and LAT040 to assess the potential release of intact *gag/pol* transcripts in virions released from CD4⁺ T cells. This assay evaluates the ability of latent HIV-1 to produce infectious virus under stimulating conditions. The procedure followed a previously published method ^34^.

### Statistical analysis

HIV-1 RNA transcriptional activity in CD4-T cells was quantified as previously described ^27^. Standard curves for absolute quantification were generated using serial dilutions of HIV-1 plasmid controls in GraphPad Prism v10 (GraphPad Software). CA HIV-1 short RNA transcripts were measured using the Double-R assay, and values were log-transformed where appropriate to approximate normality.

Continuous variables were assessed for distribution prior to analysis. Between-group comparisons were performed using non-parametric Mann–Whitney U tests for two-group comparisons (e.g., presence vs absence of detectable DRMs, or DRM class strata). For comparisons involving more than two groups (e.g., number of ART drug classes with detected DRMs), ordinal groupings were also evaluated using Mann–Whitney U tests where appropriate, given limited sample sizes and non-Gaussian distributions.

Associations between CA HIV-1 short RNA transcript levels and neuroimaging outcomes, including NAA in FWM, posterior PCC, and caudate nucleus, were assessed using Pearson correlation for linear relationships. Where appropriate, Spearman rank correlation was also considered for sensitivity analyses of non-parametric associations.

Biomarker associations with neuroaxonal integrity outcomes were assessed across systemic inflammatory markers, monocyte/macrophage activation markers, vascular risk markers, and clinical neurocognitive status (including HAND classification). Given the exploratory nature of the study, no formal adjustment for multiple comparisons was applied; instead, consistency of findings across brain regions and effect size were considered in interpretation.

For drug resistance mutation (DRM) and premature stop codon (PSC) analyses, categorical comparisons between groups were performed using Mann–Whitney U tests or Fisher’s exact tests where appropriate. All tests were two-tailed, and statistical significance was defined as p < 0.05.

## Acknowledgments

We thank the participants for their time and involvement in the study. We also acknowledge Positive Life NSW/NAPWHA for assisting with the nationwide recruitment drive. We gratefully acknowledge trial coordination by Fiona Kilkenny, Sarah Barney, and John Ng; MRI coordination and voxel placement for MRS by Kirsten Moffat; and PBMC biobanking by Kate Merlin, Sara Evans, Bertha Fsadni, Maeve Hodgson, Kevin Tran, and Ethan Lewis.

## Funding

This research was partly funded by a St Vincent’s Clinic Foundation Research Grant and an AMR Translational Research Grant to K.S., with additional support from NHMRC Grant ID1105808 to B.J.B.

### Database

Complete biomarker measurements and associated datasets will be made available through the repository described in the final published article.

## Author Contributions

K.S. conceived the project. K.S., H.O., and Y.I. designed the experiments and analyses. A.L., E.Y., and K.S. conducted the experiments. A.L., E.Y., Y.I., and H.O. performed the nanopore data analysis. T.I. contributed reagents for the short transcript analysis. M.M., I.W.D., and K.B. provided essential support and procedures for long transcript analysis using nanopore sequencing. K.S., J.Z., H.O., Y.I., and B.J.B. wrote the manuscript. L.A.C. reviewed and analyzed the 1H-MRS data and collected participants’ clinical characteristics, including HAND assessments. H.O. and Y.I. reviewed, analyzed, and generated all nanopore-based sequencing datasets. H.O. reviewed and confirmed the molecular analyses. B.J.B. reviewed and confirmed the clinical relevance of the findings. All authors reviewed and approved the final manuscript.

---

**Supplementary Figure 1.**
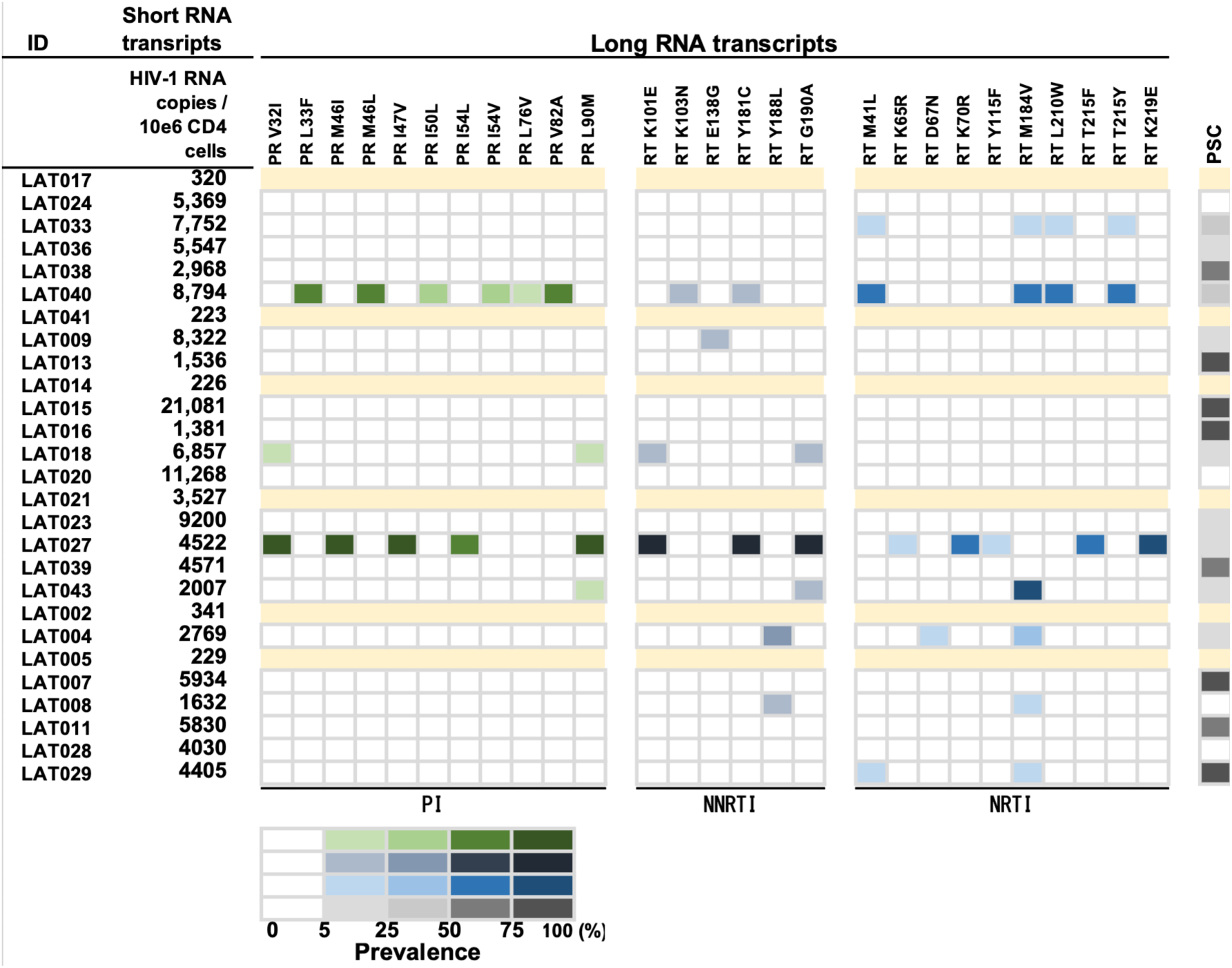
Analysis of short and long HIV-1 RNA transcripts in the hidden viral reservoir of 27 individuals. Detection of short and long HIV-1 RNA transcripts in CD4⁺ T cells from virally suppressed individuals. Long transcripts were undetectable in 6 participants (highlighted in light yellow). In the remaining 21 individuals, long transcripts were successfully amplified and analyzed for DRMs. For each sample, the within-sample prevalence of major DRMs in the *pol* region is shown, excluding sequences with premature stop codons (PSCs). Prevalence is color-coded according to the scale shown in the bar legend. The proportion of sequences with PSCs in *pol* is also indicated for each individual.

**Supplementary Figure 2.**
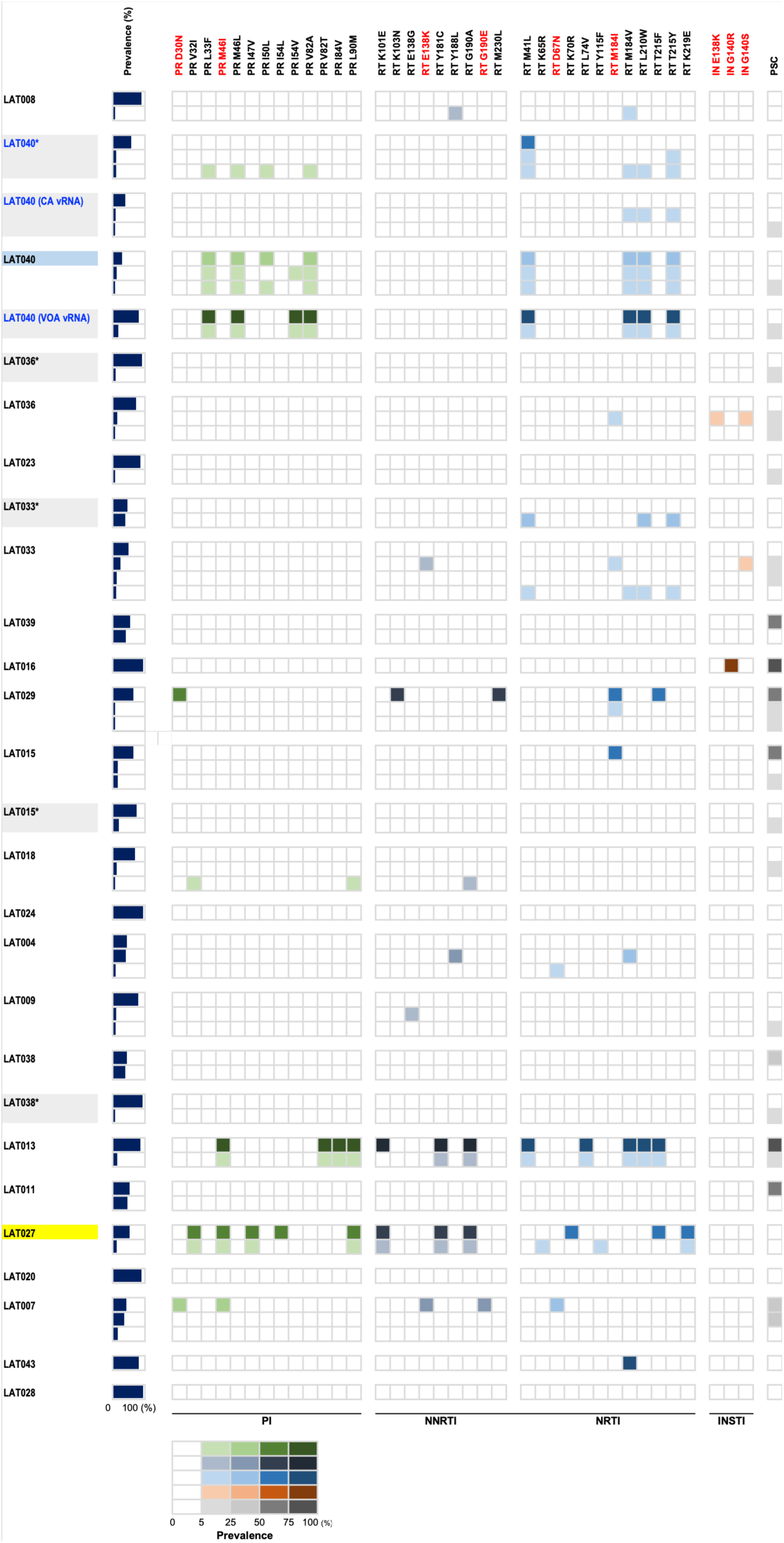
Linkage analysis of major DRMs and PSCs. Within-sample prevalence of genotypes composed of co-existent major DRMs and/or PSCs within pol. Each genotype is defined as a unique combination pattern of major DRMs in the presence or absence of co-existent PSC at any position within pol. The labels of the DRMs that could arise from APOBEC3 editing are colored in red. Prevalence of each genotype in the respective sample is color-coded according to the bottom bars in the middle part and is shown as a navy bar graph. Rare genotypes that occupied at <5% of population for each sample were omitted. The genotypes bearing DRMs that were shared among different samples within a patient are highlighted with colored arrows on the right side of the table. Samples from the same individual—including additional CA-vRNA or VOA-vRNA sequences obtained at later time points (18 or 24 months) are denoted by asterisks (*) on a light gray-shaded background. *Note for LAT040: Linkage analysis revealed that PI- and NRTI-related DRMs co-existed within the same viral genome in LAT040. However, linkage analysis for NNRTI-related DRMs was inconclusive due to their low prevalence (<5% cutoff), reflecting limited detectability*.

**Supplementary Figure 3.**
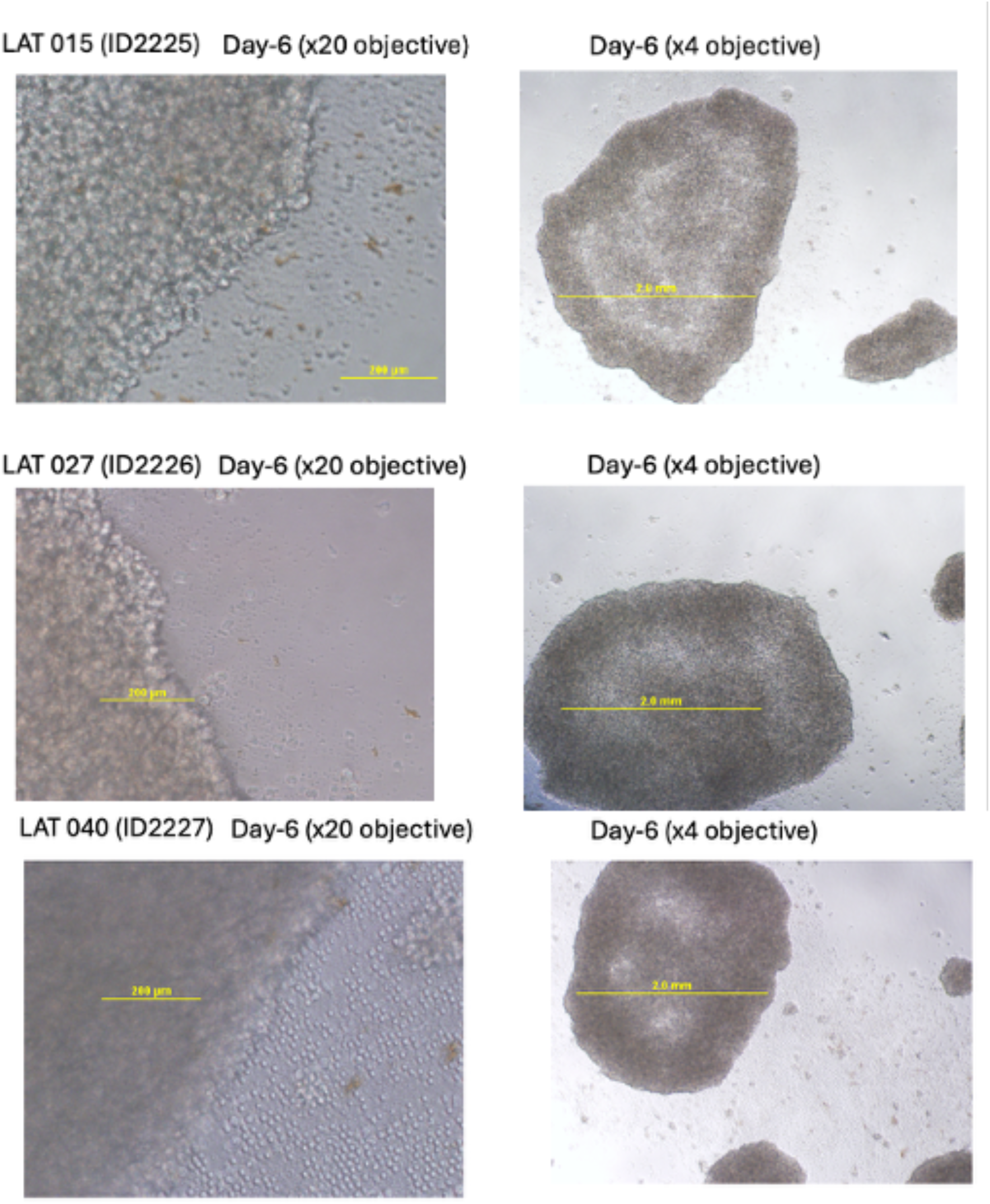
Microscopy of purified CD4⁺ T cells after 6 days in culture. Purified CD4⁺ T cells were cultured for 6 days in IL-2–containing medium in the presence of a potent T-cell activator cocktail comprising anti-CD3, anti-CD28, and anti-CD2 monoclonal antibodies (anti-CD3/CD28/CD2 activator) 26. Representative images show cell morphology and clustering under stimulation conditions.

**Supplementary Figure 4.**
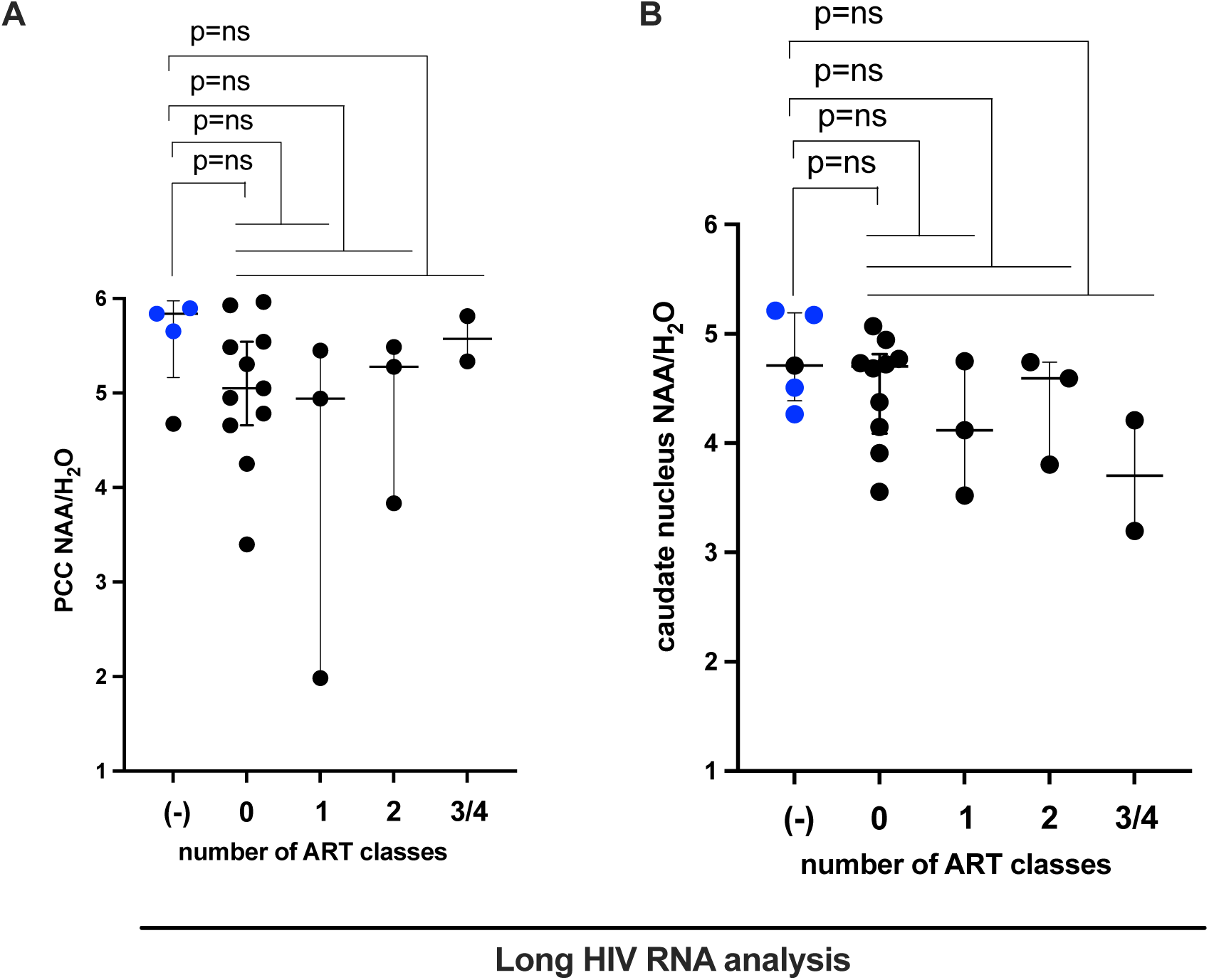
Impact of HIV-1 long RNA transcripts on neuroaxonal integrity. **(A)** Association pf PCC NAA levels in FWM with absence or presence of the Reservoir DRMs (Blue colored individuals are identical as in Figure 1C). **(B)** Association pf caudate nucleus NAA levels in FWM with absence or presence of the Reservoir DRMs (Blue colored individuals are identical as in Figure 1C).

**Supplementary Table 1.**
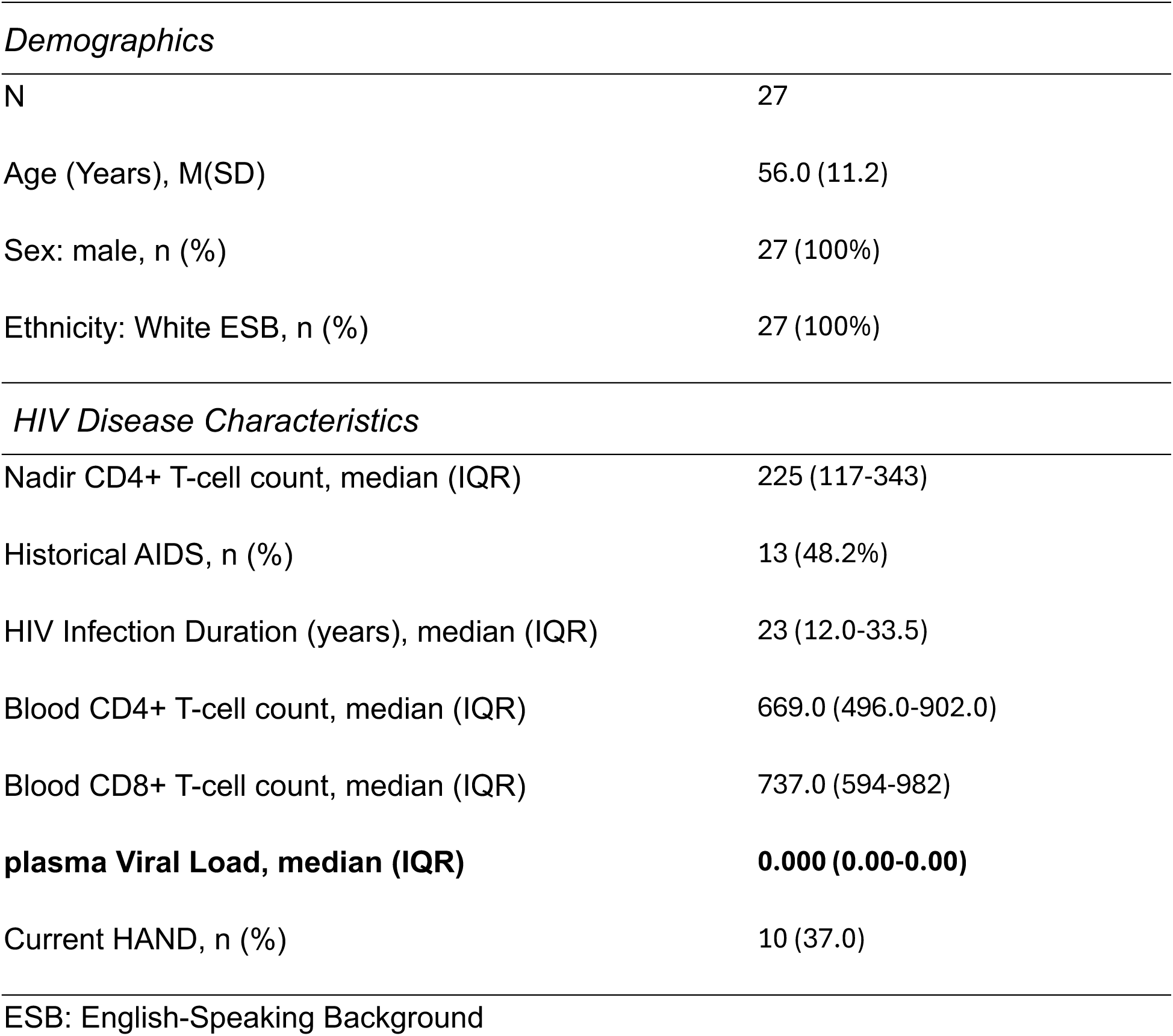
Cohort Demographics and Disease Characteristics.

|  |  |
| --- | --- |
| *Demographics* |  |
| N | 27 |
| Age (Years), M(SD) | 56.0 (11.2) |
| Sex: male, n (%) | 27 (100%) |
| Ethnicity: White ESB, n (%) | 27 (100%) |
| *HIV Disease Characteristics* |  |
| Nadir CD4+ T-cell count, median (IQR) | 225 (117-343) |
| Historical AIDS, n (%) | 13 (48.2%) |
| HIV Infection Duration (years), median (IQR) | 23 (12.0-33.5) |
| Blood CD4+ T-cell count, median (IQR) | 669.0 (496.0-902.0) |
| Blood CD8+ T-cell count, median (IQR) | 737.0 (594-982) |
| **plasma Viral Load, median (IQR)** | **0.000 (0.00-0.00)** |
| Current HAND, n (%) | 10 (37.0) |
| ESB: English-Speaking Background |  |

## References

1. Cysique, L.A., Moffat, K., Moore, D.M., Lane, T.A., Davies, N.W., Carr, A., Brew, B.J., and Rae, C. (2013). HIV, vascular and aging injuries in the brain of clinically stable HIV-infected adults: a (1)H MRS study. PLoS One 8, e61738. 10.1371/journal.pone.0061738.

2. McArthur, J.C., Brew, B.J., and Nath, A. (2005). Neurological complications of HIV infection. Lancet Neurol 4, 543–555. 10.1016/S1474-4422(05)70165-4.

3. Jamal Eddine, J., Angelovich, T.A., Zhou, J., Byrnes, S.J., Tumpach, C., Saraya, N., Chalmers, E., Shepherd, R.A., Tan, A., Marinis, S., et al. (2024). HIV transcription persists in the brain of virally suppressed people with HIV. PLoS Pathog 20, e1012446. 10.1371/journal.ppat.1012446.

4. Rosenthal, J., and Tyor, W. (2019). Aging, comorbidities, and the importance of finding biomarkers for HIV-associated neurocognitive disorders. Journal of neurovirology 25, 673–685. 10.1007/s13365-019-00735-0.

5. Lanman, T., Letendre, S., Ma, Q., Bang, A., and Ellis, R. (2021). CNS Neurotoxicity of Antiretrovirals. Journal of neuroimmune pharmacology : the official journal of the Society on NeuroImmune Pharmacology 16, 130–143. 10.1007/s11481-019-09886-7.

6. Moschopoulos, C.D., Alford, K., Antoniadou, A., and Vera, J.H. (2024). Cognitive impairment in people living with HIV: mechanisms, controversies, and future perspectives. Trends Mol Med 30, 1076–1089. 10.1016/j.molmed.2024.06.005.

7. Debuka, A., and Chakravarty, J. (2025). Neurological manifestations and neuropharmacology of HIV/AIDS. In Neuropsychiatric Complications of HIV, pp. 195–211. 10.1016/b978-0-12-818851-4.00002-2.

8. Churchill, M., and Nath, A. (2013). Where does HIV hide? A focus on the central nervous system. Curr Opin HIV AIDS 8, 165–169. 10.1097/COH.0b013e32835fc601.

9. Angelovich, T.A., Cochrane, C.R., Zhou, J., Tumpach, C., Byrnes, S.J., Jamal Eddine, J., Waring, E., Busman-Sahay, K., Deleage, C., Jenkins, T.A., et al. (2023). Regional Analysis of Intact and Defective HIV Proviruses in the Brain of Viremic and Virally Suppressed People with HIV. Ann Neurol 94, 798–802. 10.1002/ana.26750.

10. Hiener, B., Horsburgh, B.A., Eden, J.S., Barton, K., Schlub, T.E., Lee, E., von Stockenstrom, S., Odevall, L., Milush, J.M., Liegler, T., et al. (2017). Identification of Genetically Intact HIV-1 Proviruses in Specific CD4(+) T Cells from Effectively Treated Participants. Cell Rep 21, 813–822. 10.1016/j.celrep.2017.09.081.

11. Wang, Z., Simonetti, F.R., Siliciano, R.F., and Laird, G.M. (2018). Measuring replication competent HIV-1: advances and challenges in defining the latent reservoir. Retrovirology 15, 21. 10.1186/s12977-018-0404-7.

12. Berkhout, B., and de Ronde, A. (2004). APOBEC3G versus reverse transcriptase in the generation of HIV-1 drug-resistance mutations. AIDS 18, 1861–1863.

13. Piantadosi, A., Humes, D., Chohan, B., McClelland, R.S., and Overbaugh, J. (2009). Analysis of the percentage of human immunodeficiency virus type 1 sequences that are hypermutated and markers of disease progression in a longitudinal cohort, including one individual with a partially defective Vif. J Virol 83, 7805–7814. 10.1128/JVI.00280-09.

14. Brodin, J., Zanini, F., Thebo, L., Lanz, C., Bratt, G., Neher, R.A., and Albert, J. (2016). Establishment and stability of the latent HIV-1 DNA reservoir. Elife 5. 10.7554/eLife.18889.

15. Bruner, K.M., Murray, A.J., Pollack, R.A., Soliman, M.G., Laskey, S.B., Capoferri, A.A., Lai, J., Strain, M.C., Lada, S.M., Hoh, R., et al. (2016). Defective proviruses rapidly accumulate during acute HIV-1 infection. Nat Med 22, 1043–1049. 10.1038/nm.4156.

16. Chun, T.W., Davey, R.T., Jr., Engel, D., Lane, H.C., and Fauci, A.S. (1999). Re-emergence of HIV after stopping therapy. Nature 401, 874–875. 10.1038/44755.

17. Galvao-Lima, L.J., Zambuzi, F.A., Soares, L.S., Fontanari, C., Meireles, A.F.G., Brauer, V.S., Faccioli, L.H., Gama, L., Figueiredo, L.T.M., Bou-Habib, D.C., and Frantz, F.G. (2022). HIV-1 Gag and Vpr impair the inflammasome activation and contribute to the establishment of chronic infection in human primary macrophages. Molecular immunology 148, 68–80. 10.1016/j.molimm.2022.04.018.

18. Richard, J., Prevost, J., Bourassa, C., Brassard, N., Boutin, M., Benlarbi, M., Goyette, G., Medjahed, H., Gendron-Lepage, G., Gaudette, F., et al. (2023). Temsavir blocks the immunomodulatory activities of HIV-1 soluble gp120. Cell Chem Biol 30, 540–552 e546. 10.1016/j.chembiol.2023.03.003.

19. Baiyegunhi, O.O., Mthembu, K., Reuschl, A.K., Ojwach, D., Farinre, O., Maimela, M., Balinda, S., Price, M., Bunders, M.J., Altfeld, M., et al. (2025). HIV-1 Gag-protease-driven replicative capacity influences T-cell metabolism, cytokine induction, and viral cell-to-cell spread. mBio 16, e0356524. 10.1128/mbio.03565-24.

20. Solis, M., Nakhaei, P., Jalalirad, M., Lacoste, J., Douville, R., Arguello, M., Zhao, T., Laughrea, M., Wainberg, M.A., and Hiscott, J. (2011). RIG-I-mediated antiviral signaling is inhibited in HIV-1 infection by a protease-mediated sequestration of RIG-I. J Virol 85, 1224–1236. 10.1128/JVI.01635-10.

21. Berg, R.K., Melchjorsen, J., Rintahaka, J., Diget, E., Soby, S., Horan, K.A., Gorelick, R.J., Matikainen, S., Larsen, C.S., Ostergaard, L., et al. (2012). Genomic HIV RNA induces innate immune responses through RIG-I-dependent sensing of secondary-structured RNA. PLoS One 7, e29291. 10.1371/journal.pone.0029291.

22. Telwatte, S., Moron-Lopez, S., Aran, D., Kim, P., Hsieh, C., Joshi, S., Montano, M., Greene, W.C., Butte, A.J., Wong, J.K., and Yukl, S.A. (2019). Heterogeneity in HIV and cellular transcription profiles in cell line models of latent and productive infection: implications for HIV latency. Retrovirology 16, 32. 10.1186/s12977-019-0494-x.

23. Olson, R.M., Gornalusse, G., Whitmore, L.S., Newhouse, D., Tisoncik-Go, J., Smith, E., Ochsenbauer, C., Hladik, F., and Gale, M., Jr. (2022). Innate immune regulation in HIV latency models. Retrovirology 19, 15. 10.1186/s12977-022-00599-z.

24. Anderson, A.M., Munoz-Moreno, J.A., McClernon, D.R., Ellis, R.J., Cookson, D., Clifford, D.B., Collier, A.C., Gelman, B.B., Marra, C.M., McArthur, J.C., et al. (2017). Prevalence and Correlates of Persistent HIV-1 RNA in Cerebrospinal Fluid During Antiretroviral Therapy. J Infect Dis 215, 105–113. 10.1093/infdis/jiw505.

25. Spudich, S., Robertson, K.R., Bosch, R.J., Gandhi, R.T., Cyktor, J.C., Mar, H., Macatangay, B.J., Lalama, C.M., Rinaldo, C., Collier, A.C., et al. (2019). Persistent HIV-infected cells in cerebrospinal fluid are associated with poorer neurocognitive performance. J Clin Invest 129, 3339–3346. 10.1172/JCI127413.

26. Chen, W., Berkhout, B., and Pasternak, A.O. (2025). Phenotyping Viral Reservoirs to Reveal HIV-1 Hiding Places. Curr HIV/AIDS Rep 22, 15. 10.1007/s11904-025-00723-6.

27. Suzuki, K., Zaunders, J., Gates, T.M., Levert, A., Butterly, S., Liu, Z., Ishida, T., Palmer, S., Rae, C.D., Juge, L., et al. (2022). Elevation of cell-associated HIV-1 transcripts in CSF CD4+ T cells, despite effective antiretroviral therapy, is linked to brain injury. Proc Natl Acad Sci U S A 119, e2210584119. 10.1073/pnas.2210584119.

28. Swanton, C., McGranahan, N., Starrett, G.J., and Harris, R.S. (2015). APOBEC Enzymes: Mutagenic Fuel for Cancer Evolution and Heterogeneity. Cancer discovery 5, 704–712. 10.1158/2159-8290.CD-15-0344.

29. Jakobsdottir, G.M., Brewer, D.S., Cooper, C., Green, C., and Wedge, D.C. (2022). APOBEC3 mutational signatures are associated with extensive and diverse genomic instability across multiple tumour types. BMC Biol 20, 117. 10.1186/s12915-022-01316-0.

30. Lu, H., Lu, Z., Wang, Y., Chen, M., Li, G., and Wang, X. (2025). APOBEC in breast cancer: a dual player in tumor evolution and therapeutic response. Front Mol Biosci 12, 1604313. 10.3389/fmolb.2025.1604313.

31. Derache, A., Shin, H.S., Balamane, M., White, E., Israelski, D., Klausner, J.D., Freeman, A.H., and Katzenstein, D. (2015). HIV drug resistance mutations in proviral DNA from a community treatment program. PLoS One 10, e0117430. 10.1371/journal.pone.0117430.

32. Pasternak, A.O., and Berkhout, B. (2018). What do we measure when we measure cell-associated HIV RNA. Retrovirology 15, 13. 10.1186/s12977-018-0397-2.

33. Vignoles, M., Andrade, V., Noguera, M., Brander, C., Mavian, C., Salemi, M., Paredes, R., Sharkey, M., and Stevenson, M. (2021). Persistent HIV-1 transcription in CD4(+) T cells from ART-suppressed individuals can originate from biologically competent proviruses. J Virus Erad 7, 100053. 10.1016/j.jve.2021.100053.

34. Suzuki, K., Levert, A., Yeung, J., Starr, M., Cameron, J., Williams, R., Rismanto, N., Stark, T., Druery, D., Prasad, S., et al. (2021). HIV-1 viral blips are associated with repeated and increasingly high levels of cell-associated HIV-1 RNA transcriptional activity. AIDS. 10.1097/QAD.0000000000003001.

35. Dyer, W.B., Suzuki, K., Levert, A., Starr, M., Lloyd, A.R., and Zaunders, J.J. (2024). Preservation of functionality, immunophenotype, and recovery of HIV RNA from PBMCs cryopreserved for more than 20 years. Front Immunol 15, 1382711. 10.3389/fimmu.2024.1382711.

36. Suzuki, K., Gold, L., Levert, A., Butterly, S., Yoo, E., Ishida, T., Zaunders, J., Cysique, L.A., and Brew, B.J. (2025). Persistent cell-associated HIV-1 RNA in virally suppressed individuals on INSTI-based ART. Journal of Virus Eradication 11. 10.1016/j.jve.2025.100609.

37. Suzuki, K., Levert, A., Yoo, E., Cysique, L.A., Ishida, T., Olsen, N., Brew, B.J., and Zaunders, J. (2025). Higher levels of cell-associated HIV-1 RNA transcripts predict risk of viral blips: implications for clinical management and cure research. AIDS 39, 1300–1302. 10.1097/QAD.0000000000004191.

38. Sarkhouh, H., and Chehadeh, W. (2021). CODEHOP-Mediated PCR Improves HIV-1 Genotyping and Detection of Variants by MinION Sequencing. Microbiol Spectr 9, e0143221. 10.1128/Spectrum.01432-21.

39. Mori, M., Ode, H., Kubota, M., Nakata, Y., Kasahara, T., Shigemi, U., Okazaki, R., Matsuda, M., Matsuoka, K., Sugimoto, A., et al. (2022). Nanopore Sequencing for Characterization of HIV-1 Recombinant Forms. Microbiol Spectr 10, e0150722. 10.1128/spectrum.01507-22.

40. Delaney, K.E., Ngobeni, T., Woods, C.K., Gordijn, C., Claassen, M., Parikh, U., Harrigan, P.R., and van Zyl, G.U. (2023). Nano-RECall provides an integrated pipeline for HIV-1 drug resistance testing from Oxford Nanopore sequence data. Trop Med Int Health 28, 186–193. 10.1111/tmi.13851.

41. Ode, H., Matsuda, M., Shigemi, U., Mori, M., Yamamura, Y., Nakata, Y., Okazaki, R., Kubota, M., Setoyama, Y., Imahashi, M., et al. (2024). Population-based nanopore sequencing of the HIV-1 pangenome to identify drug resistance mutations. Sci Rep 14, 12099. 10.1038/s41598-024-63054-3.

42. Karn, J. (1999). Tackling Tat. J Mol Biol 293, 235–254.

43. Karn, J., and Stoltzfus, C.M. (2012). Transcriptional and posttranscriptional regulation of HIV-1 gene expression. Cold Spring Harb Perspect Med 2, a006916. 10.1101/cshperspect.a006916.

44. Mbonye, U., and Karn, J. (2017). The Molecular Basis for Human Immunodeficiency Virus Latency. Annu Rev Virol 4, 261–285. 10.1146/annurev-virology-101416-041646.

45. Brady, J., and Kashanchi, F. (2005). Tat gets the “green” light on transcription initiation. Retrovirology 2, 69. 10.1186/1742-4690-2-69.

46. Mbonye, U., Kizito, F., and Karn, J. (2023). New insights into transcription elongation control of HIV-1 latency and rebound. Trends Immunol 44, 60–71. 10.1016/j.it.2022.11.003.

47. Pollard, V.W., and Malim, M.H. (1998). The HIV-1 Rev protein. Annu Rev Microbiol 52, 491–532. 10.1146/annurev.micro.52.1.491.

48. Emerman, M., and Malim, M.H. (1998). HIV-1 regulatory/accessory genes: keys to unraveling viral and host cell biology. Science 280, 1880–1884. 10.1126/science.280.5371.1880.

49. Cullen, B.R., and Malim, M.H. (1991). The HIV-1 Rev protein: prototype of a novel class of eukaryotic post-transcriptional regulators. Trends Biochem Sci 16, 346–350. 10.1016/0968-0004(91)90141-h.

50. Coffin, J.M., Hughes, S.H., and Varmus, H.E. (1997). Retroviruses. Retroviruses, 205–261 Cold Spring Harbor Laboratory Press, Cold Spring Harbor, NY.

51. Aksenova, M.V., Silvers, J.M., Aksenov, M.Y., Nath, A., Ray, P.D., Mactutus, C.F., and Booze, R.M. (2006). HIV-1 Tat neurotoxicity in primary cultures of rat midbrain fetal neurons: changes in dopamine transporter binding and immunoreactivity. Neurosci Lett 395, 235–239. 10.1016/j.neulet.2005.10.095.

52. Zhang, T., Ding, H., An, M., Wang, X., Tian, W., Zhao, B., and Han, X. (2020). Factors associated with high-risk low-level viremia leading to virologic failure: 16-year retrospective study of a Chinese antiretroviral therapy cohort. BMC Infect Dis 20, 147. 10.1186/s12879-020-4837-y.

53. Ajasin, D., and Eugenin, E.A. (2020). HIV-1 Tat: Role in Bystander Toxicity. Front Cell Infect Microbiol 10, 61. 10.3389/fcimb.2020.00061.

54. Muvenda, T., Williams, A.A., and Williams, M.E. (2024). Transactivator of Transcription (Tat)-Induced Neuroinflammation as a Key Pathway in Neuronal Dysfunction: A Scoping Review. Mol Neurobiol 61, 9320–9346. 10.1007/s12035-024-04173-w.

55. Coffin, J.M., Hughes, S.H., and Varmus, H.E. (1997). The Interactions of Retroviruses and their Hosts. In Retroviruses, J.M. Coffin, S.H. Hughes, and H.E. Varmus, eds.

56. Bansal, A.K., Mactutus, C.F., Nath, A., Maragos, W., Hauser, K.F., and Booze, R.M. (2000). Neurotoxicity of HIV-1 proteins gp120 and Tat in the rat striatum. Brain Res 879, 42–49. 10.1016/s0006-8993(00)02725-6.

57. Potter, M.C., Figuera-Losada, M., Rojas, C., and Slusher, B.S. (2013). Targeting the glutamatergic system for the treatment of HIV-associated neurocognitive disorders. Journal of neuroimmune pharmacology : the official journal of the Society on NeuroImmune Pharmacology 8, 594–607. 10.1007/s11481-013-9442-z.

58. Huiting, E.D., Gittens, K., Justement, J.S., Shi, V., Blazkova, J., Benko, E., Kovacs, C., Wender, P.A., Moir, S., Sneller, M.C., et al. (2019). Impact of Treatment Interruption on HIV Reservoirs and Lymphocyte Subsets in Individuals Who Initiated Antiretroviral Therapy During the Early Phase of Infection. J Infect Dis 220, 270–274. 10.1093/infdis/jiz100.

59. Cohn, L.B., Chomont, N., and Deeks, S.G. (2020). The Biology of the HIV-1 Latent Reservoir and Implications for Cure Strategies. Cell Host Microbe 27, 519–530. 10.1016/j.chom.2020.03.014.

60. De Bellis, A., Willemsen, M.S., Guzzetta, G., van Sighem, A., Romijnders, K., Reiss, P., Schim van der Loeff, M.F., van de Wijgert, J., Nijhuis, M., Kretzschmar, M.E.E., and Rozhnova, G. (2025). Model-based evaluation of the impact of a potential HIV cure on HIV transmission dynamics. Nature communications 16, 3527. 10.1038/s41467-025-58657-x.

61. Deng, Z., Yan, H., Lambotte, O., Moog, C., and Su, B. (2025). HIV controllers: hope for a functional cure. Front Immunol 16, 1540932. 10.3389/fimmu.2025.1540932.

62. Kuhn, L. (2025). Stepping stones to cure in children with HIV. Curr Opin HIV AIDS 20, 247–248. 10.1097/COH.0000000000000925.

63. Lanzafame, M., Mori, G., and Vento, S. (2025). Advances in HIV Treatment: Long-Acting Antiretrovirals and the Path Toward a Cure. Biomedicines 13. 10.3390/biomedicines13020493.

64. Xu, P., Yuan, D., Moog, C., and Su, B. (2025). Efforts toward achieving the goal of ending AIDS by 2030: from antiretroviral drugs to HIV vaccine and cure research. Sci China Life Sci. 10.1007/s11427-024-2840-4.

65. Luth, T., Schaake, S., Grunewald, A., May, P., Trinh, J., and Weissensteiner, H. (2022). Benchmarking Low-Frequency Variant Calling With Long-Read Data on Mitochondrial DNA. Front Genet 13, 887644. 10.3389/fgene.2022.887644.

66. Vandenhende, M.A., Bellecave, P., Recordon-Pinson, P., Reigadas, S., Bidet, Y., Bruyand, M., Bonnet, F., Lazaro, E., Neau, D., Fleury, H., et al. (2014). Prevalence and evolution of low frequency HIV drug resistance mutations detected by ultra deep sequencing in patients experiencing first line antiretroviral therapy failure. PLoS One 9, e86771. 10.1371/journal.pone.0086771.

67. Price, M.N., Dehal, P.S., and Arkin, A.P. (2010). FastTree 2--approximately maximum-likelihood trees for large alignments. PLoS One 5, e9490. 10.1371/journal.pone.0009490.

